# The unresolved phylogenomic tree of butterflies and moths (Lepidoptera): assessing the potential causes and consequences

**DOI:** 10.1101/2021.04.09.439156

**Authors:** Jadranka Rota, Victoria Twort, Andrea Chiocchio, Carlos Peña, Christopher W. Wheat, Lauri Kaila, Niklas Wahlberg

## Abstract

The field of molecular phylogenetics is being revolutionised with next-generation sequencing technologies making it possible to sequence large numbers of genomes for non-model organisms ushering us into the era of phylogenomics. The current challenge is no longer how to get enough data, but rather how to analyse the data and how to assess the support for the inferred phylogeny. We focus on one of the largest animal groups on the planet – butterflies and moths (order Lepidoptera). We clearly demonstrate that there are unresolved issues in the inferred phylogenetic relationships of the major lineages, despite several recent phylogenomic studies of the group. We assess the potential causes and consequences of the conflicting phylogenetic hypotheses. With a dataset consisting of 331 protein-coding genes and the alignment length over 290 000 base pairs, including 200 taxa representing 81% of lepidopteran superfamilies, we compare phylogenetic hypotheses inferred from amino acid and nucleotide alignments. The resulting two phylogenies are discordant, especially with respect to the placement of the superfamily Gelechioidea, which is likely due to compositional bias of both the nucleotide and amino acid sequences. With a series of analyses, we dissect our dataset and demonstrate that there is sufficient phylogenetic signal to resolve much of the lepidopteran tree of life. Overall, the results from the nucleotide alignment are more robust to the various perturbations of the data that we carried out. However, the lack of support for much of the backbone within Ditrysia makes the current butterfly and moth tree of life still unresolved. We conclude that taxon sampling remains an issue even in phylogenomic analyses, and recommend that poorly sampled highly diverse groups, such as Gelechioidea in Lepidoptera, should receive extra attention in the future.

## Introduction

The advent of next-generation sequencing technologies has led to a revolution in the field of molecular phylogenetics. The era of phylogenomics has arrived with new genomes for non-model organisms being sequenced at an unprecedented scale. Now researchers must consider how to best use the acquired data to infer phylogenetic relationships while acknowledging the ongoing challenges that remain. In phylogenomic studies, protein-coding genes are a commonly used source of phylogenetic information, in particular when sequencing transcriptomes. This raises an additional question, how we should analyse the data, as amino acids or nucleotides, as well as whether the different codon positions in protein-coding genes should be treated equally. Our reading of literature suggests that it is often thought that amino acid sequences are appropriate for deeper phylogenetic problems, while DNA nucleotide sequences are more informative at shallower levels. However, at intermediate levels it has been noted that there can be conflicts between these two types of data (e.g. Simmons et al. 2002; Zwick et al. 2012; Gillung et al. 2018; Vasilikopoulos et al. 2019). Often, when there is such conflict, it seems to be more common to present the results from amino acid analyses as the preferred hypothesis (see our literature review for examples; supplementary file 1). Here we assess this preference for amino acid level analyses by investigating whether choosing amino acids based inferences over nucleotides produces unbiased insights or skews our understanding of the tree of life.

Perhaps the largest problem inherent to the era of phylogenomics is the sheer size of datasets, which typically consist of hundreds to thousands of genes. In such cases we are often restricted to bioinformatics methods for determining primary homologies of sequences (orthology and alignments), which commonly results in a situation unthinkable in the pre-phylogenomics era, that researchers do not actually know what their data look like. In shifting to a phylogenomics scale, we necessarily have black-boxed our data and must largely trust our bioinformatics tools to determine orthology and align sequences correctly. Despite rigorous procedures to retain only alignments passing some threshold of scores designed to assess the quality of the alignments, alignment problems can persist with regard to homologies (Wong et al. 2008; Philippe et al. 2011; Di Franco et al. 2019).

Based on the number of described species, butterflies and moths (order Lepidoptera) are one of the largest groups of macro-organisms on Earth. Molecular data have shed much light on the phylogenetic relationships of the families in this large group (Regier et al. 2009; Mutanen et al. 2010; Regier et al. 2013; Heikkilä et al. 2015), and the latest datasets based on transcriptomes and target enrichment appear to be converging on a consensus of the deeper relationships within the order (Bazinet et al. 2013; Kawahara and Breinholt 2014; Bazinet et al. 2017; Kawahara et al. 2019; Mayer et al. 2021). These datasets are based on single copy orthologous protein-coding genes, ranging from 741 (Bazinet et al. 2013) to 1753 in Mayer et al. (2021), 2098 (Kawahara et al. 2019), 2696 (Kawahara and Breinholt 2014), and 2948 genes (Breinholt et al. 2018). The phylogenomic studies support many of the surprising results obtained in previous Sanger sequencing based studies, such as the position of Pyraloidea as the sister to the so-called macro-moths (Macroheterocera), and butterflies as a lineage unrelated to the macro-moths.

On the other hand, there are clear conflicts between these phylogenomic studies. The most striking conflict is within Apoditrysia – a large clade including the well-known lepidopterans such as butterflies, macro-moths, leaf-rollers, snout moths, and many others. One of the apoditrysian groups, plume moths (Pterophoroidea), is found to be in a clade with Thyridoidea and Gelechioidea in Bazinet et al. (2013), Kawahara et al. (2019), and Mayer et al. (2021), but in a clade with leaf-rollers (Tortricoidea) and Urodoidea in Kawahara and Breinholt (2014) and Breinholt et al. (2018). The position of other apoditrysian groups, such as the Cossoidea–Zygaenoidea complex, is also unstable (Mitter et al. 2017). Similarly, some conflicts exist in the relationships of the macro-moths, such as inchworms and relatives (Geometroidea) being sister to owlet moths (Noctuoidea) with full support in Kawahara and Breinholt (2014), while in Bazinet et al. (2013), Kawahara et al. (2019), and Mayer et al. (2021) Geometroidea is sister to silk moths and relatives (Bombycoidea + Lasiocampoidea). Some of these discrepancies could perhaps be explained by very low taxon sampling in the earliest phylogenomic studies (e.g. Bazinet et al. 2013).

What becomes obvious at closer inspection is that in some of these studies the presented results are based on nucleotide alignments (e.g. Bazinet et al. 2013; Kawahara and Breinholt 2014), while in others they are based on amino acid alignments (e.g. Kawahara et al. 2019). For some of the conflicting branches, node support values are low, but for others they are high. It is clear that the backbone of the clade Ditrysia (which contains 98% of the extant 160,000 described species; van Nieukerken et al. 2011) remains poorly supported, even when datasets comprise thousands of genes and more than two million base pairs (bp), as well as representatives of most superfamilies. Here we attempt to uncover some of the reasons behind the observed discordance in results from different phylogenomic studies. We produced a set of 331 manually vetted genes for 195 species representative of 34 of 42 currently recognized lepidopteran superfamilies, and we used this dataset to test four hypotheses explaining the possible reasons for the lack of concordance among the various published phylogenies.

Hypothesis 1 (H1): the lack of concordance results from insufficient signal in the data. We test this hypothesis through a close examination of branch support in trees based on nucleotide and amino acid datasets and the comparison of the topology with other published studies. If we find that the overall topology is well supported and concordant with topologies derived from other datasets, we can falsify this hypothesis and conclude that it is not the lack of phylogenetic signal that is behind the discordant results.

Hypothesis 2 (H2): the lack of concordance results from gene tree/species tree issues, in other words, the evolutionary history of genes differs from that of the species. If it turns out that the differences in topology mainly stem from analyses of different datasets and not amino acid/nucleotide analyses of the same data, this could imply that we are dealing with gene tree versus species tree issues and need to conduct analyses employing the multispecies coalescent model.

Hypothesis 3 (H3): the lack of concordance results from the type of data analysed, where analyses of nucleotide alignments give one topology and those of amino acid alignments give another topology. We test this hypothesis by comparing results from analyses on amino acid and nucleotide alignments in our study as well as other similar studies. The focus here is on the relationships within Apoditrysia – a large clade that is highly unstable.

Hypothesis 4 (H4): the lack of concordance results from compositional bias. Compositional bias (also known as compositional heterogeneity) refers to different nucleotide or amino acid composition in some taxa and is known to potentially mislead phylogenetic inference (Jermiin et al. 2004). If there is support for H3, the question arises what is behind the difference in topologies resulting from the amino acid and nucleotide alignments. A possible explanation is systematic bias arising from compositional bias. Compositional bias could affect one or both types of data. We test this hypothesis by first carrying out tests for compositional bias and then proceeding to exclude taxa/genes with a high compositional bias to ascertain how their exclusion influences the topology.

We test the numbered hypotheses (H1–4), and we propose other topics, such as model misspecification and a potential rapid radiation, as future avenues for exploration. Finally, we discuss the implications of our findings for our understanding of the Lepidoptera phylogeny, as well as for the field of phylogenomics in general.

## Material and methods

### Literature review

We carried out a literature review to establish whether amino acid or nucleotide alignments are more commonly analysed in insect phylogenomic studies and whether the authors present results from both types of analyses equally. We examined all of the papers published on lepidopteran phylogenomics as well as papers published on phylogenomics of other insect groups from the following journals: BMC Evolutionary Biology, Molecular Phylogenetics and Evolution, Systematic Biology, and Systematic Entomology in 2019 and 2020. We excluded studies that had only mitochondrial DNA data even if the authors presented them as phylogenomic studies.

### Taxon sampling

Our dataset includes 195 ingroup species representing 34 of 42 currently recognized lepidopteran superfamilies and 72 of 137 families (van Nieukerken et al. 2011; Kristensen et al. 2015; Rajaei et al. 2015; Regier et al. 2015a; Kaila et al. 2020). Of the 34 sampled superfamilies, 17 are monotypic (i.e. including only one family). For six superfamilies we sampled all of the families included in them (Drepanoidea, Geometroidea, Neopseustoidea, Nepticuloidea, Papilionoidea, and Pyraloidea). For eight superfamilies we sampled more than 1/3 of the included families, while three superfamilies are relatively poorly sampled (Gelechioidea, Hepialoidea, and Yponomeutoidea). The species included are listed in Table S1. Additionally, we used five species of the lepidopteran sister group Trichoptera as outgroups.

### Genomes and transcriptomes

Genomes (a total of eight) were downloaded from the NCBI Assembly database (Kitts et al. 2016), while transcriptomes (a total of 192) were downloaded as either raw reads from the Sequence Read Archive (SRA; Leinonen et al. 2011) or as assemblies if available. The bulk of the transcriptomes are from three major phylogenomic studies on Lepidoptera (Bazinet et al. 2013; Kawahara and Breinholt 2014; Bazinet et al. 2017). A number of Lepidoptera transcriptomes were also available from other studies, including a few critical taxa not sampled in the three major studies (see Table S1).

For specimens without assembly, the raw reads from SRA were processed as follows. Reads were trimmed to remove adapter sequences and low-quality regions (Q < 30 and Q < 20 for Illumina and 454 data, respectively) were removed using Cutadapt 1.4.1 (Martin 2011) (minimum read length 50 bp). Homopolymer stretches were removed with Prinseq v 0.20.4 (Schmieder and Edwards 2011). Transcriptomes were *de novo* assembled using Trinity 2.0.6 (Grabherr et al. 2011; Haas et al. 2013), set to default parameters with a minimum contig length of 100 bp and a minimum kmer coverage of 5. Redundancy among the resulting transcripts was reduced using CD-HIT-EST v3.1.1 with a similarity threshold of 95% (Li and Godzik 2006).

### Selecting genes

Single copy orthologous protein coding genes were compiled from the *Bombyx mori* genome (GCA_000151625) based on 1478 genes reported by Misof et al. (2014). In addition, we also included all ribosomal protein genes, mitochondrial protein genes, as well as the standard Sanger sequenced genes used in Lepidoptera systematics (Wahlberg and Wheat 2008; Regier et al. 2009), for a total of 1551 protein coding genes (Supplementary material).

Gene screening was carried out on an initial set of 119 taxa, including eight whole genomes and 111 transcriptomes available at that time (2015, Table S1). Assemblies were queried against the gene set using a tBlastn (v2.2.28, Altschul et al. 1990) approach (1e-5 cut-off).

The output of the BLAST search (coordinates of BLAST hits within each transcriptome) was then used to extract the DNA sequences from each assembly using a set of Python scripts written by CP (https://github.com/carlosp420/PyPhyloGenomics).

The resulting output files were scanned manually using two criteria. First, we wanted to find the commonly expressed genes occurring in a wide variety of transcriptomes that were produced for a variety of purposes and encompassed different life stages and tissue types. For this criterion, we kept only those genes that were found in at least 80% of the target transcriptomes.

Second, the retained set of genes were scanned and aligned manually based on amino acid translations. Only those genes that could be unambiguously aligned manually based on the amino acid translation were kept. If there was any ambiguity in alignment, the gene was discarded. In five cases, a large gene had an ambiguously aligned region in the middle (highlighted in Table S2). In these cases, the gene was divided into two or three unambiguously aligned regions, and the ambiguous regions were discarded. Some genes were fragmented in some transcriptomes, with missing segments. These genes were compiled manually for maximum representation.

After screening, a set of 331 genes was chosen as a representative gene set, divided into 337 loci. These loci were used as query sequences for additional BLAST searches.

An SRA search (May 2017) identified a further 81 transcriptomes to be added to the existing dataset, mainly from Bazinet et al. (2017). The 331 gene set was screened using a bidirectional blast approach, either tBlastn or Blastx (1e-5 cut-off). DNA sequences from each assembly were extracted using the same python script mentioned above. The resulting sequences were translated to amino using TranslatorX (Abascal et al. 2010) and aligned to the existing alignments using MAFFT 7.266 (Katoh et al. 2005; Katoh and Standley 2013), using the add fragments and auto options to preserve existing gaps in the alignment and choose the most appropriate alignment strategy, respectively. Alignments were manually screened and then converted back to nucleotide alignments with pal2nal v14 (Suyama et al. 2006). Aligned DNA sequences were curated and maintained in VoSeq (Peña and Malm 2012), from which the various datasets were generated. The final dataset was 97 206 amino acids or 194 410 bp (NT12) long.

### Phylogenetic analyses

Initially we analysed the nucleotide data with all codon positions included, but this resulted in a highly anomalous tree compared to the accepted lepidopteran relationships (result not shown). As the third codon positions are rarely phylogenetically informative for older divergences because of the high substitution rate resulting in rapid saturation, we proceeded to analyse the data with only first and second codon positions included (NT12_all). We also analysed amino acid sequences (AA_all), as well as the NT12 data applying the so-called degen1 coding (Regier et al. 2010; Zwick et al. 2012), which codes for only non-synonymous changes (NT12_degen1). These datasets included all 200 taxa and 331 genes.

All analyses were done in the maximum likelihood framework using IQ-TREE v.1.6.8 (Nguyen et al. 2015) on the CIPRES server (Miller et al. 2010). Altogether, we analysed 21 different datasets (Table 1). Of the analysed datasets, nine were amino acid (AA) datasets, one was analysed with degen1 coding, nine were nucleotide datasets including first and second codon positions (NT12), and one included all three codon positions (Papilionoidea, NT123).

**Table 1.** An overview of the 19 datasets analysed in this study. NT12 – nucleotide dataset including first and second codon positions; AA – amino acid dataset; degen1 – using degenerative coding for nucleotides that makes synonymous changes invisible (see text for more details).

| Dataset | Dataset description |
| --- | --- |
| NT12, AA_all | 200 taxa, 331 genes |
| NT12_degen1 | 200 taxa, 331 genes, degen1 coding |
| NT12, AA_no_highGC | 200 taxa, 281 genes (50 high-GC-content genes excluded) |
| NT12, AA_noGele | 193 taxa (all 7 Gelechioidea excluded), 331 genes |
| NT12, AA_noTorts | 192 taxa (all 8 Tortricidae excluded), 331 genes |
| NT12, AA_minus2Gele | 198 taxa (2 gelechioids with low compositional bias excluded), 331 genes |
| NT12, AA_minus3Gele | 197 taxa (3 gelechioids with high compositional bias excluded), 331 genes |
| NT12, AA_minus10 | 200 taxa, 321 genes (10 genes with highest support for NT/AA all topology excluded) |
| NT12, AA_minus20 | 200 taxa, 311 genes (20 genes with highest support for NT/AA all topology excluded) |
| NT12, AA_minus50 | 200 taxa, 281 genes (50 genes with highest support for NT/AA all topology excluded) |
| NT12, Papilionoidea | 40 taxa, 331 genes |
| NT123, Papilionoidea | 40 taxa, 331 genes |

The data were analysed as a single partition (degen1) or partitioned by gene (all other datasets). For each dataset, the command TESTNEW was used, i.e. the ModelFinding algorithm (Kalyaanamoorthy et al. 2017) with rate heterogeneity modelled through the probability distribution free model. The best models were then used to estimate the maximum likelihood topology with 100 randomly built parsimony trees as starting trees (command -t RANDOM). Robustness of the phylogenetic hypothesis was assessed with 1000 replicates of ultrafast bootstraps (UFB) (Hoang et al. 2018) and 1000 replicates of the SH-like approximate likelihood ratio test (SH) (Guindon et al. 2010). We follow the recommendation that clades with UFB≥95 and SH≥80 can be considered as well supported. For comparison of overall branch support across different trees from the various analyses that we carried out, we used the average UFB values from all of the nodes in the backbone of the tree up to the clade Macroheterocera (aUFB).

In addition to using SH, UFB, and aUFB to assess branch support, we also calculated gene and site concordance factors (sCF) in IQ-tree (Minh et al. 2020). Gene concordance factors (gCF) are the percentage of single-locus trees that are consistent with a particular branch in the reference tree, while site concordance factors (sCF) express the percentage of sites that support the same resolution of a quartet around a particular node as in the reference tree. For these reasons, gCF values range 0–100% whereas sCF values are around 33% when the data have no signal for resolving a quartet in question, and are otherwise higher. We calculated these factors on the NT12_all dataset.

Because these values were lower than what we were expecting, we made a more extensive exploration of these factors in a subset of our data, including only the taxa belonging to the superfamily Papilionoidea, known as butterflies. The phylogeny of butterflies is very well known and has changed very little with significant increases in the amount of sequence (Espeland et al. 2018, 207 species, 352 genes) or significant increases in taxon sampling (Chazot et al. 2019, 994 species, 10 genes). These two most recent studies, which are based on different datasets, are highly concordant with each other, as well as with the earlier efforts (Heikkilä et al. 2012). The topology for this group in our NT12_all and AA_all datasets is fully concordant with both of these studies and we use it to better understand the different values of gCF and sCF. As the CF values for the butterflies were also low in the NT12 dataset, we reasoned that this could be because 3^rd^ codon positions would actually contain phylogenetic signal at this shallower level in the tree. Papilionoidea are estimated to be ca. 108 My old, (Chazot et al. 2019), while Lepidoptera are about 300 My old (Kawahara et al. 2019) so we repeated these analyses on the NT123 dataset for Papilionoidea.

### Hypothesis testing

H1: the lack of concordance results from insufficient signal in the data. We compared tree topologies resulting from our various analyses to published results by focusing on the following 10 key relationships that have consistently received high support: 1) Micropterigidae + all others; 2) Angiospermivora; 3) Glossata; 4) Heteroneura; 5) Eulepidoptera; 6) Euheteroneura; 7) Ditrysia; 8) Apoditrysia; 9) Pyraloidea + Macroheterocera; 10) Macroheterocera. These clades (marked in Fig. 1) span the whole breadth of the lepidopteran tree of life and if they are all present in all of the resulting topologies, we can conclude that there is enough phylogenetic signal in our dataset to infer these relationships. We compare our results to the two most recent published studies, which have similar taxon sampling to ours: Kawahara et al. (2019) (2098 genes) and Mayer et al. (2021) (1753 genes). As Heterobathmioidea are not sampled in Mayer et al. (2021) and the support for Angiospermivora can therefore not be assessed, we focus on the remaining nine clades in this set of comparisons. When evaluating strength of branch support, we consider values of UFB=>95 or bootstrap=>90 to denote strong support.

**Figure 1.**
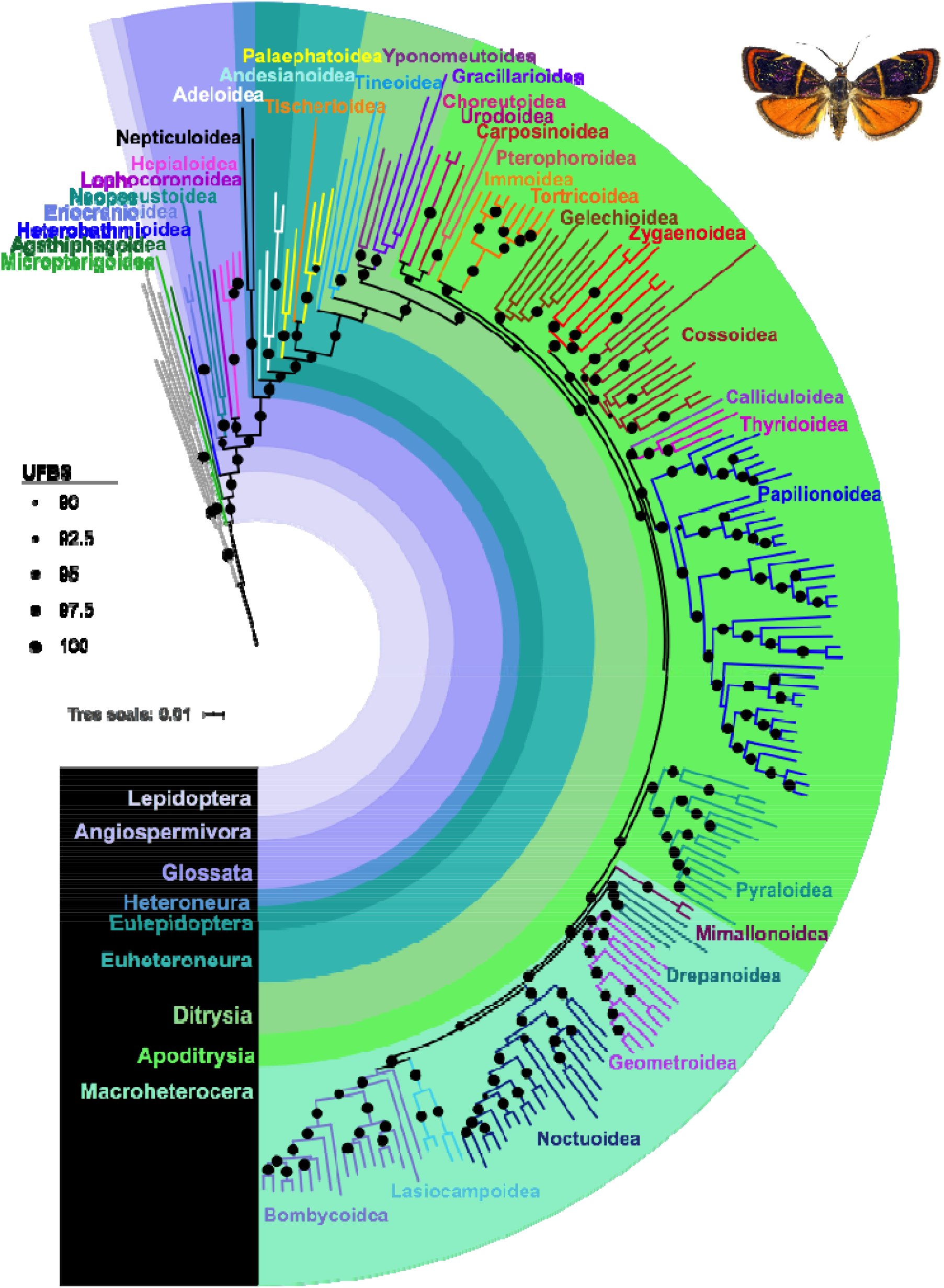
A phylogenetic tree from an analysis of 331 gene fragments from 300 taxa (including five caddisfly outgroups) representing 34 out of 41 superfamilies of butterflies and moths inferred using maximum likelihood on a nucleotide alignment with 3rd codon positions excluded (dataset NT12_all) in IQ-Tree. Dots on nodes represent ultrafast bootstraps (UFBS). Major clades that are well supported are marked as well as all superfamilies.

Hypothesis 2 (H2): the topological discordance is the result of gene trees being different from species trees. To test for this, we performed multispecies coalescent analyses using the method implemented in ASTRAL 5.6.3 (Zhang et al. 2018). At first, we filtered each gene alignment to remove sequences with 100% missing data using the Alignment_assesment.py and Alignment_refinement.py scripts in Phyton from Portik et al. (2016). Then, for both the NT12_all and AA_all dataset, we estimated single gene trees using RAxML 8.2.10 (Stamatakis 2014), with the GTRGAMMA model of DNA evolution and 100 bootstrap replicates for branch support. Finally, we ran ASTRAL using the default options. We compared the ASTRAL trees with those from concatenated alignments both for our datasets and the two most recent Lepidoptera phylogenomic studies (Kawahara et al. 2019; Mayer et al. 2021).

Hypothesis 3 (H3): the lack of concordance results from the type of data analysed, where analyses of nucleotide alignments give one topology and those of amino acid alignments give another topology. We test this hypothesis by comparing tree topologies focusing on Apoditrysia, which is the most unstable part of the lepidopteran tree of life. We score trees from different analyses for the presence of the following taxa in the same clade, even if the clade might include additional members: 1) Choreutoidea + Urodoidea; 2) Immoidea + Tortricoidea; 3) Pterophoroidea + Carposinoidea + Urodoidea; 4) Pterophoroidea + Carposinoidea; 5) Cossoidea + Zygaenoidea; 6) Papilionoidea + Calliduloidea + Thyridoidea; 7) Papilionoidea + (Pyraloidea + Macroheterocera); 8) Gelechioidea + Calliduloidea + Thyridoidea; 9) Pterophoroidea + Gelechioidea + Calliduloidea + Thyridoidea; 10) Noctuoidea + (Lasiocampoidea + Bombycoidea); 11) Geometroidea + (Lasiocampoidea + Bombycoidea); 12) (Geometroidea + Noctuoidea) + (Bombycoidea + Lasiocampoidea). The following topologies were compared: NT12_all, NT12_degen1, AA_all (this study; Table 1); Bazinet et al. (2013) NT123 degen1; Kawahara & Breinholt (2014) NT123, AA; Breinholt et al. (2018) dataset 3 (NT12 degen1, 2696 genes); Kawahara et al. (2019) NT degen1, AA; and Mayer et al. (2021) NT12, AA. The overlap in taxon sampling is high between the Kawahara et al. (2019), Mayer et al. (2021) and our study, making these comparisons easy. Kawahara and Breinholt (2014) are missing a few of the key taxa, rendering some of the comparisons impossible.

As there is significant overlap in data between all of these studies, they are not independent from each other. Bazinet et al. (2013), and the Kawahara and Breinholt (2014) studies have very large overlap. The Mayer et al. (2021) dataset has a large overlap with the Kawahara et al. (2019) since the same data were mined for their hybrid enrichment (Breinholt et al. 2018). Out of 331 genes in our study, 180 are also present in the Kawahara et al. (2019) dataset, and 56 are shared with the Mayer et al. (2021) dataset.

Hypothesis 4 (H4): systematic bias in phylogenetic inference is arising from compositional bias. To evaluate the influence of compositional heterogeneity and the possible impact of sequences whose evolution violated the assumptions of global stationarity, reversibility, and homogeneity, we conducted a pairwise sequence comparison as implemented in SymTest version 2.0.47 (Jermiin et al. 2004; Ababneh et al. 2006). We applied the Bowker’s matched-pairs tests of symmetry (Bowker 1948) on both the amino acid and the nucleotide dataset without 3^rd^ codon positions, using the default values for gene window and step-size, and we generated the heat maps based on the inferred p-values for each comparison. Since we discovered high compositional heterogeneity especially in Tortricoidea and Gelechiodea (see Results), we sorted all genes by their GC content in Tortricoidea and removed 50 genes with the highest GC content (Table S2). We carried out phylogeny inference on the resulting dataset (no_high_GC) in the same way as for the complete datasets. Since sampled gelechioids and tortricids had genes with the highest relative GC content (see Results), we wanted to test the effect of removing these groups on the topology. In these analyses, all gelechioids (no_Gele) or all tortricids (no_Torts) were excluded.

Additionally, we excluded several but not all gelechioid taxa from some of the analyses because the Bowker’s test results showed that there was a high amount of compositional heterogeneity within Gelechioidea. Two of the seven sampled gelechioids (Depressariidae: *Tonica nigricostella* and Lecithoceridae: *Thubana* sp.) have a similar nucleotide composition to most of the sampled lepidopterans in this study while three of them are quite different (Depressariidae: *Psilocorsis reflexella*; Gelechiidae: *Dichomeris punctidiscella* and *Tuta absoluta*); see the Results and Discussion section for more details. Therefore, in one set of analyses we excluded *Tonica* and *Thubana* (minus2Gele) and in another set we excluded *Psilocorsis, Dichomeris*, and *Tuta* (minus3Gele) for both the AA and NT12 alignments and compared the resulting trees.

### Topology robustness and branch support

In our attempt to carry out further data exploration and to gain a better understanding of where the phylogenetic signal is coming from, we examined the strength of phylogenetic signal of different genes. Following Shen et al. (2017), we measured phylogenetic signal of every gene in both the amino acid and nucleotide alignments for the two alternative topologies (NT12_all and AA_all). We then removed from each of the alignments successively 10, 20, and 50 genes (minus_10, minus_20, and minus_50 datasets) with the strongest signal for the topology resulting from that alignment. In other words, NT12_minus10 represents a nucleotide alignment from which we removed 10 genes that had the highest phylogenetic signal for the NT12_all topology. The reasoning behind these analyses is to assess to what extent the topology is stable when genes with the strongest phylogenetic signal are removed.

## Results and Discussion

### Literature review

Searching through recently published studies (2019, 2020) including phylogenomic datasets of different insect groups, we found 23 papers in addition to the 11 phylogenomic studies published on Lepidoptera by the end of 2020, for a total of 34 studies (Table 2; see Supplementary file 1 for details and complete list of references). Thirteen of these studies were at the level of the order, seven were above the family but below order level, and 14 were at the family or lower level. At the level of the order, in 7/13 studies the analyses were carried out both on the amino acids and nucleotides and in four of them the authors noted that the results between these two types of analyses were conflicting. At the intermediate level, above family but below order, in 5/7 studies the authors did both amino acid and nucleotide analyses and four of these showed conflicts in results. At the lowest taxonomic level, in 5/14 studies the authors carried out both types of analyses and conflicting results were noted in two of these studies. Overall, at the higher taxonomic level, it was more common to see analyses of amino acid alignments (10/13 at the order level vs. 4/14 at the family or lower level). It is interesting that in some studies the authors chose to display the results from the amino acid analyses even though they had lower branch support (e.g. Kawahara et al. 2019). It appears that, as we were expecting, at the higher taxonomic levels it is more common to analyse amino acids while this is more rarely done at the family or lower level. Conflicts in results between these two types of analyses are quite common, found in 59% of these studies, thus we find it surprising that this issue has not been investigated in more detail.

**Table 2.** An overview of literature review of 34 recent studies with phylogenomic datasets of insects showing in how many studies the authors carried out which type of analyses. Thirteen studies were at the level of the order, seven above the family but below order level, and 14 at the family or lower level. AA – amino acid dataset; NT123 – nucleotide dataset with all three codon positions included; NT reduced – nucleotide dataset with first and/or third codon positions excluded; NT degen – nucleotide dataset with degen coding (see text for explanation). For studies that included both AA and NT datasets, the number of studies with conflict between the two is provided. Full reference data available in Table S2.

| Taxonomic level<br>(No. studies) | Order<br>(13 studies) | Order < Family<br>(7 studies) | ≤ Family<br>(14 studies) |
| --- | --- | --- | --- |
| AA and NT | 7 (4 w/ conflict) | 5 (4 w/ conflict) | 5 (2 w/ conflict) |
| AA | 10 | 5 | 4 |
| NT123 | 3 | 7 | 14 |
| NT reduced | 5 | 2 | 0 |
| NT degen | 4 | 2 | 1 |

### Hypothesis testing

H1: the lack of concordance results from insufficient signal in the data: The ten key clades that span the breadth and depth of the lepidopteran tree of life are strongly supported in both of our complete datasets (NT12_all, AA_all; Figs 1, S1, S2). All of them are supported in the Kawahara et al. (2019) AA dataset, and all nine that are present in the Mayer et al. (2021) dataset are supported in their analyses as well (Table 3). This result falsifies H1 as it appears that these datasets have enough phylogenetic signal to resolve the relationships in much of the lepidopteran tree of life. Overall, these relationships largely agree not only with other molecular studies (e.g. Regier et al. 2015b; Bazinet et al. 2017) but also with the morphological phylogeny of Lepidoptera (Kristensen and Skalski 1998).

**Table 3.**
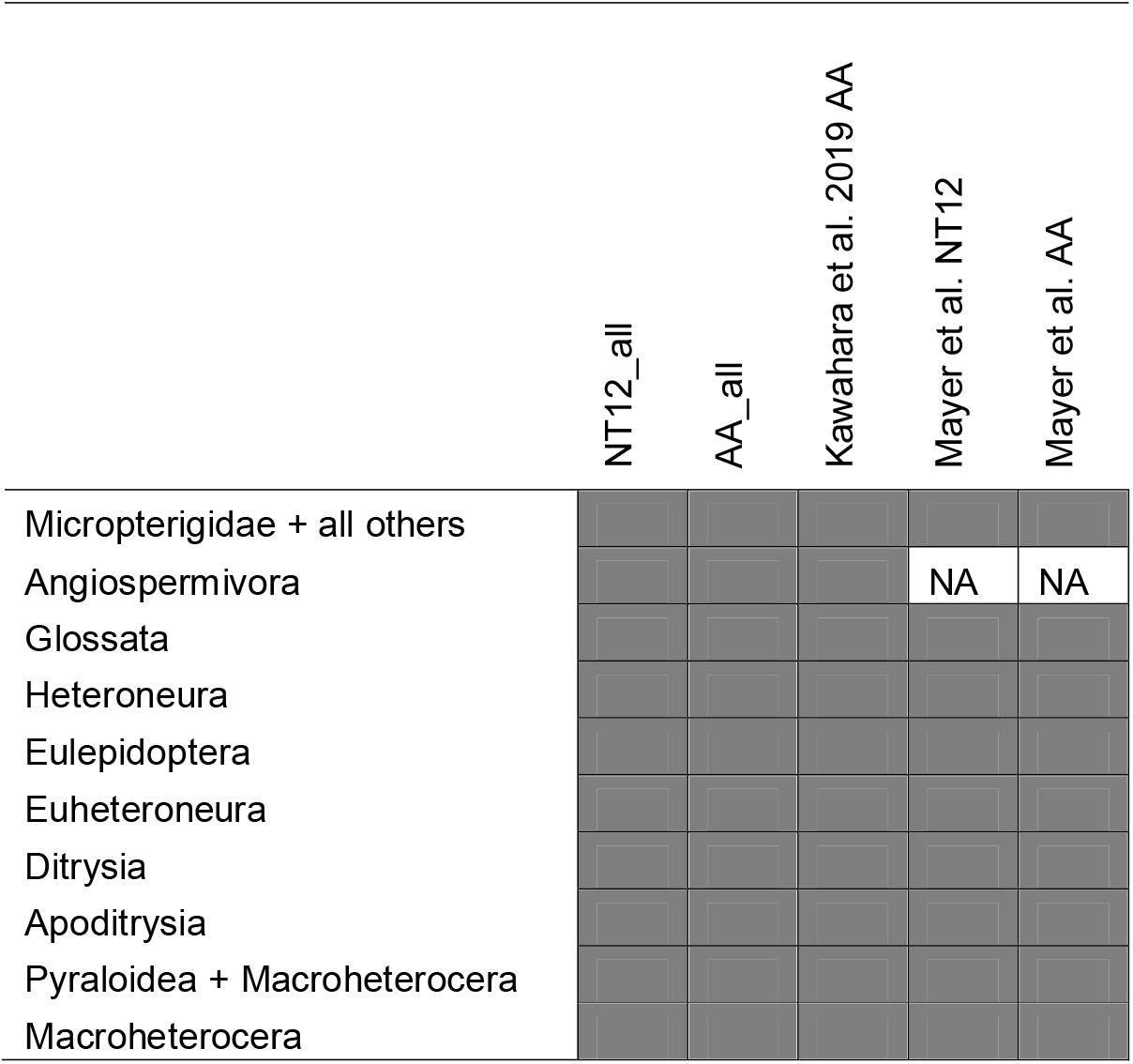
Ten key clades whose presence in topologies resulting from different datasets and analysis types was used to assess hypothesis 1 (insufficient data for resolving the lepidopteran tree of life) are listed in the rows and the datasets scored for their presence are listed in the columns. Dark grey signifies strong support and NA (not applicable) refers to taxa not being sampled in a particular dataset. All ten clades are marked in Fig. 1.

However, it can still be that there is insufficient data or signal for resolving parts of the tree, reflecting potential rapid radiations of some lineages, such as Ditrysia. The inferred internal branches among the various apoditrysian superfamilies are visibly shorter than among the non-ditrysian superfamilies, strongly suggesting that the apoditrysian lineages have experienced a rapid radiation (Figs 1, S1, S2). While superfamilies and families can be easily separated to a great extent, i.e. their monophyly is well supported, exactly how they diverged from each other is a much more difficult question to answer.

H2: the topological discordance is the result of some gene trees being different from species trees. The ASTRAL tree from the NT12 dataset (Fig. S3) is in some respects similar to the one resulting from the concatenated dataset, but in other respects it is very different. For example, some of the similarities are that Gelechioidea are recovered in the same position but with very low support (0.36), Urodoidea + Pterophoroidea are recovered as sisters (0.71) and they are together sister to Carposinoidea (0.43). On the other hand, Calliduloidea + Thyridioidea are recovered as sister to Pyraloidea + Macroheterocera (0.52); Tortricoidea are recovered as sister to all other Apoditrysia (0.94); and the topology for non-ditrysians is quite different in a number of ways. In the AA ASTRAL tree, the topology for the non-ditrysia is also in conflict with the concatenated AA tree (Fig. S4). What is interesting is that in this analysis Tortricoidea are also recovered as sister to all other Apoditrysia (0.91), but this is where similarities between the AA and NT12 ASTRAL trees for apoditrysian relationships end. Carposinoidea are recovered as sister to Pyraloidea (0.23); and Calliduloidea, Pterophoroidea, Gelechioidea, and Thyridioidea are recovered in the same part of the tree, but with Thyridoidea sister to Macroheterocera (0.35). The extremely short branches in this part of the tree make it impossible to assess the actual relationships among these groups. In Macroheterocera, Drepanoidea are recovered as paraphyletic with *Doa* (Doidae) being sister to Geometroidea (0.53).

The comparison of the Kawahara et al. (2019) AA ASTRAL tree with their concatenated tree shows that similar differences in topologies exist there as well, including an anomalous arrangement within non-ditrysians. For example, Urodoidea are recovered as sister to all Apoditrysia (0.92), Alucitoidea + Pterophoroidea (0.75) are recovered as sister to Papilionoidea (0.68), and Noctuoidea are recovered as sister to Bombycoidea + Lasiocampoidea (0.86) unlike in the analysis of the concatenated dataset where Geometroidea are sister to Bombycoidea + Lasiocampoidea. However, the placement of Gelechioidea is the same – they are recovered as sister to Calliduloidea + Thyridoidea (0.42), which are together sister to Pyraloidea + Macroheterocera (0.53). Mayer et al. (2021) also found differences between the results from ASTRAL and those from the concatenated analyses.

These observed differences between the ASTRAL trees and those resulting from analyses of concatenated datasets do not necessarily mean that there is gene tree versus species tree conflict. One weakness of an ASTRAL analysis in these cases is that the species tree is estimated from gene trees where each gene tree is based on fragments that are likely much too short to recover divergences that happened more than a hundred million years ago (Apoditrysia are estimated to be about 130 MY old; Wahlberg et al. 2013; Kawahara et al. 2019). In our dataset the average gene fragment length is 864 bp and the median, at 705 bp, is even smaller. The average gene length is even shorter in the Mayer et al. (2021) dataset (ca. 401 bp) although it is somewhat longer in the Kawahara et al. dataset (ca. 1072 bp). As the relationships among the oldest lineages (non-ditrysians) are especially problematic in the ASTRAL trees where conflict is strong not only with the concatenated phylogenomic datasets, but also with a morphological phylogeny based on a wealth of characters (Kristensen and Skalski 1998) that has largely been supported with molecular data, we deem that we cannot properly test H2, trying to infer deep-level relationships with such short gene fragments.

H3: the lack of concordance within Apoditrysia results from the type of data analysed, where analyses of nucleotide alignments give one topology and those of amino acid alignments give another topology. The examination of the presence and branch support for 12 clades within Apoditrysia provides some support for this hypothesis (Table 4). The observed pattern is somewhat mixed, with the existence of most clades being more dependent on the dataset analysed rather than the type of alignment, but some clades appear to be an exception to this, and their recovery depends on the alignment analysed. For example, Gelechioidea are recovered in a different position in our AA and NT12 datasets, and the position of Papilionoidea and the large clade including Gelechiodea and several other smaller superfamilies changes in the Mayer et al. (2021) dataset depending on the alignment analysed. Likewise, the sister group relationship between Choreutoidea and Urodoidea is also dependent on the alignment: it is recovered in our NT12_all and AA_all, but not in NT12_degen1 (Fig. S5). On the other hand, the position of Pterophoroidea is highly dependent on the dataset and this superfamily is recovered in the same clade with Urodoidea only in our datasets, regardless of the alignment analysed, whereas it is recovered in the clade with Gelechioidea, Calliduloidea and Thyridoidea in the other datasets. One of the twelve clades – Immoidea + Tortricoidea – was recovered from all of the analysed datasets that included both groups (Immidae were not sampled in Kawahara and Breinholt 2014).

**Table 4.**
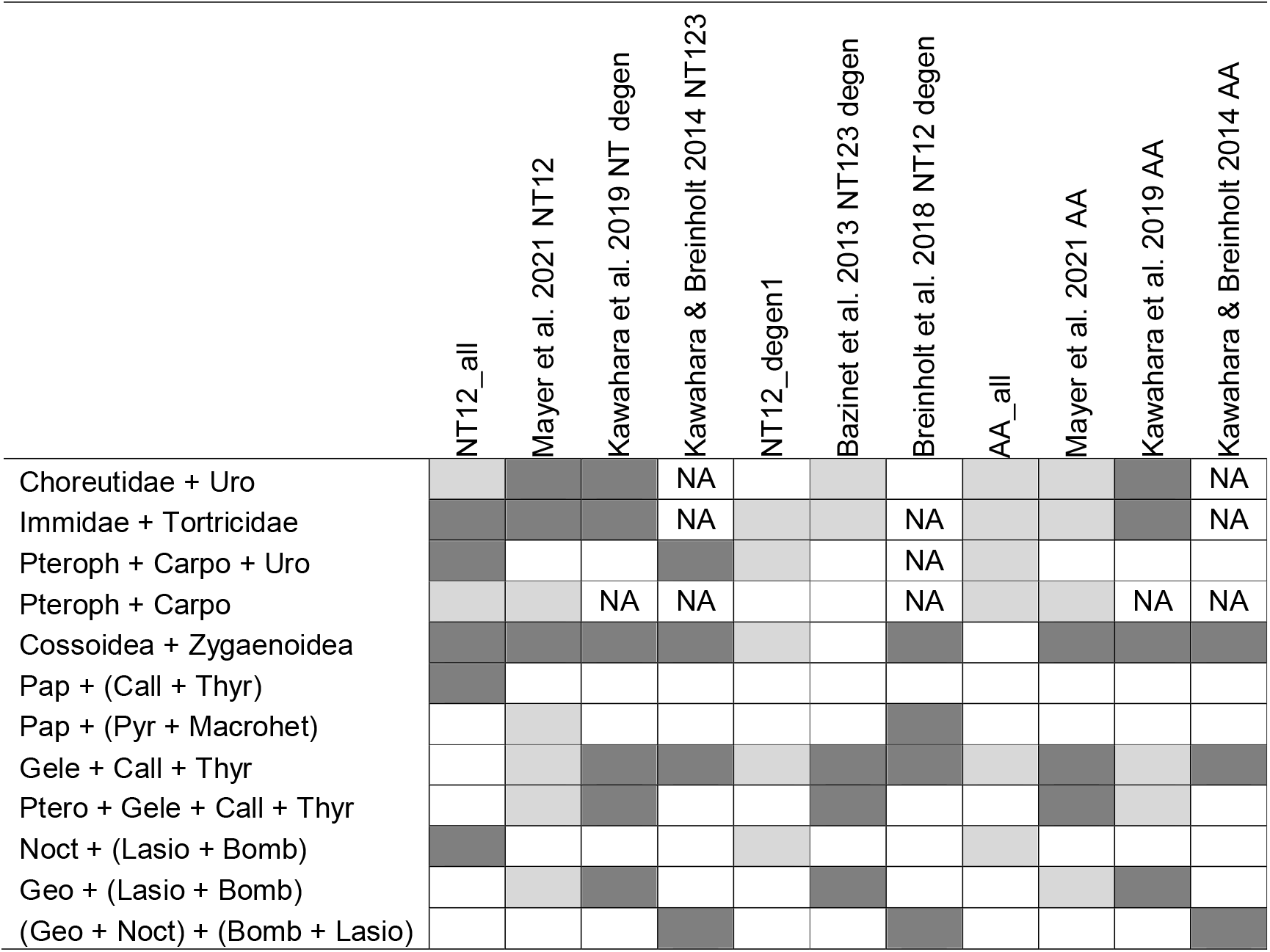
Twelve key clades within Apoditrysia whose presence in topologies resulting from different datasets and analysis types was used to assess hypothesis 2 (type of data analysed – NT vs. AA – is responsible for topological conflicts in the lepidopteran phylogeny) are listed in the rows and the datasets scored for their presence are listed in the columns. Dark grey signifies strong support, light grey signifies presence in the best tree but low support, white signifies absence in the best tree, and NA (not applicable) refers to taxa not being sampled in a particular dataset. The following abbreviations for clades/superfamilies/families are used: Bomb – Bombycoidea; Call – Calliduloidea; Carpo – Carposinoidea; Gele – Gelechioidea; Geo – Geometroidea; Pap – Papilionoidea; Lasio – Laciocampoidea; Macrohet – Macroheterocera; Noct – Noctuoidea; Pteroph – Pterophoroidea; Pyr – Pyraloidea; Thyr – Thyridoidea; Uro – Urodoidea. All the groups are shown in Fig. 1.

Possible explanations for different placement of groups in analyses of the same data are systematic bias arising for example from compositional bias (H4) or other types of model misspecifications. Possible explanations for differences in topology from different sets of genes are gene tree versus species tree issues (H2), which we addressed above. Another possible explanation for the instability of the results is relatively low taxon sampling across all of these studies. Each new study tends to include some previously unsampled groups but with the sampling efforts of the most recent studies (Kawahara et al. 2019 81% superfamilies and 54% families represented; Mayer et al. 2021 81% superfamilies, 58% families; and our study 81% superfamilies, 53% families) we are still far from having representatives of all lepidopteran families, and low sampling of some groups is probably one of the reasons behind the instability of the results.

H4: systematic bias in phylogenetic inference is arising from compositional bias. We tested if compositional heterogeneity among taxa affected our phylogenetic inference. It appears that sequences in the nucleotide dataset are unlikely to have evolved under global stationary, time-reversible, and homogeneous conditions since more than 90% of Bowker’s tests significantly rejected global symmetry. In contrast, most of the Bowker’s tests performed on the amino acid dataset were not significant, with the main exception of sequences from Tineoidea, Tortricoidea, and Gelechioidea (Figs S6–7).

Removing Tortricoidea, one of the groups with a high GC content, had no effect on the topology with respect to the placement of Gelechioidea in either AA or NT12 datasets (Figs 2c–d, S8–9) although the placement of the smaller apoditrysian superfamilies such as Choreutoidea (in both AA and NT12) and Carposinoidea (in NT12) did change compared to the AA_all and NT12_all topologies (Fig. 2a-d). Removing Gelechioidea had no effect on the topology of the NT12 dataset (Figs 2e, S10), but it did change the AA topology (Figs 2f, S11) making it very similar to the NT12_all topology as Calliduloidea + Thyridoidea were now recovered as sister to Papilionoidea albeit with no support (UFB 78, SH 67.5). However, when the three gelechioids with high compositional bias were removed, we recovered the remaining Gelechioidea as sister to Pyraloidea + Macroheterocera although not together with Calliduloidea and Thyridoidea in the NT12 topology (Fig. S12; but note that there is no support for this placement). In other words, Gelechioidea were recovered in a very different position from the NT12_all analysis. Moreover, when the two gelechioids with low compositional bias were removed, the remaining gelechioids were recovered in the exact same position as in the NT12_all analysis (Fig. S13), suggesting that the compositional bias misleads the phylogenetic inference. At the same time, removing either of these groups of gelechioids had no effect on the AA topology (Figs S14–15), but removing the three with a high bias resulted in higher UFB for the sister relationship between Gelechioidea and Calliduloidea + Thyridoidea (92 in AA_all, 98 in minus3Gele_AA; SH was 100 with and without these taxa).

**Figure 2.**
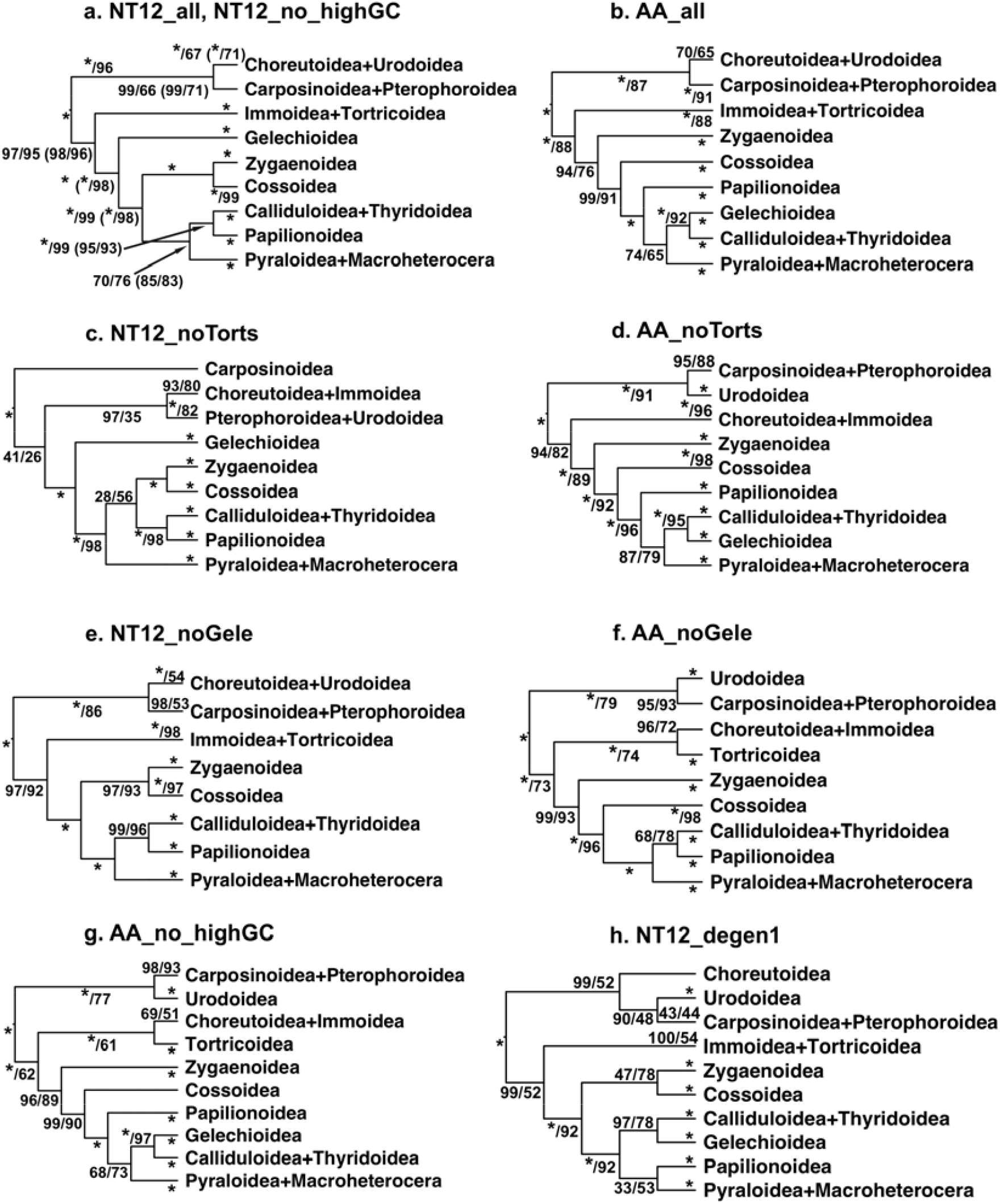
An overview of phylogenetic relationships among apoditrysian Lepidoptera from maximum likelihood analyses of different datasets. Full dataset descriptions are available in Table 1; NT12 – nucleotide alignment including 1st and 2nd codon positions; AA – amino acid alignment; degen1 – degenerative coding of nucleotides; all – all taxa and genes included; no_highGC – 50 genes with high GC content excluded; noTorts – Tortricoidea excluded; noGele – Gelechioidea excluded. Numbers and characters on branches represent branch support as SH-like approximate likelihood ratio test / ultrafast bootstrap (UFBS), with “*” denoting 100. In panel a. branch support is shown first for the NT12_all topology and in parentheses for the NT12_no_highGC topology when the numbers are different.

Removing genes with high GC content while keeping all 200 taxa in the analysis had no effect on the topology recovered from the NT12 dataset (Figs 2a, S16), while in the AA tree, Choreutoidea + Immoidea were recovered as sister to Tortricoidea (no support; Figs 2g, S17), unlike in the AA_all tree in which Choreutoidea are sister to Urodoidea (also no support) and Immoidea are alone sister to Tortricoidea (high SH support but low UFB; Fig. 2b, S2). The position of Gelechioidea did not change in either analysis.

Given that the position of Immoidea tends to change between different analyses, we examined more closely their Bowker’s test results as well. It is noteworthy that the one sampled representative of Immoidea has a very similar compositional bias to Tortricoidea. This begs the question of whether these two lineages share base composition because of their common ancestry or rather, this bias in their nucleotide composition is driving their inferred sister relationship. This is an especially relevant question as Immoidea are considered to belong to Obtectomera based on their morphology, while Tortricoidea are not (Kristensen and Skalski 1998). Obtectomera is a clade within Apoditrysia, defined based on the presence of the pupal stage with appendages fused with the body as well as immobile abdominal segments 1–4 (Kristensen and Skalski 1998). However, in Immoidea the pupal shell is weakly sclerotized and the intersegmental mobility is ambiguous, making its placement in Obtectomera questionable (Heikkilä et al. 2015). The homology of this character has already been questioned by its presence in Gelechioidea, which were placed in non-obtectomeran Apoditrysia by Minet (1991) and Kristensen & Skalski (1998) (but see Rammert 1994; Kaila 2004). A thorough discussion of this character and its phylogenetic implications is given in Heikkilä et al. (2015).

Taken all together, this strongly suggests that it is compositional bias that is driving at least some of these relationships. Removing all of the Gelechioidea results in almost the same topology for both the AA_all and NT12_all and perhaps even more tellingly, removing gelechioids with a high and low compositional bias results in two very different positions for the remaining Gelechioidea in the NT12 tree. Gelechioidea are heavily undersampled in phylogenomic studies. The clade is highly diverse with ca. 18,000 described species (van Nieukerken et al. 2011; Heikkilä et al. 2014), yet only representatives of five out of 21 families have genomic level data available (Kawahara et al. 2019, Mayer et al. 2021).

### Topology robustness and branch support

Amino acid alignments were analysed as a single partition or partitioned by gene, with no effects on the topology. Likewise, results from analysing the NT12 data with various partitioning strategies had no effect. The AA_all and NT12_all datasets yielded almost identical and a very well supported topology for the non-ditrysian groups as well as for the early branching Ditrysia. Relationships among the non-ditrysian lineages were congruent among all of the datasets, and largely congruent with Bazinet et al. (2017). However, relationships in the clade Apoditrysia were highly sensitive to data perturbations. In particular, the position of the superfamily Gelechioidea is highly dependent on the form of the data, with the NT12 datasets placing it as sister to a large clade including Zygaenoidea, Cossoidea, Papilionoidea, Pyraloidea and Macroheterocera (Figs 1, S1), while the AA datasets tend to place Gelechioidea as sister to Thyridoidea + Calliduloidea, with this clade sister to Pyraloidea + Macroheterocera (Fig. S2).

As has been demonstrated in other studies using concordance factors (Chan et al. 2020; Kallal et al. 2020; Minh et al. 2020; van Elst et al. 2021), many branches with perfect UFB and SH support have quite low CF values (Fig. S18). In the topology from the NT12_all dataset, these values are extremely low especially for the backbone nodes (average gCF=7%). The branches in the backbone are the shortest branches in the tree, and short branches have been shown to have low CF values (Chan et al. 2020; Minh et al. 2020). For most branches in this topology, the gCF values are much lower than the sCF values, suggesting that the sites that support this topology are scattered across the different genes. What we find surprising is that even for the branches leading to the lepidopteran superfamilies, almost all of which are well delineated with morphological synapomorphies and whose monophyly has been repeatedly supported in various molecular datasets, both the gCF and sCF values are low. When we exclude four superfamilies whose sampled members are close relatives and for which we obtained very high CF values (gCF>90%, sCF>80%) (members from the same family: Hapialoidea or from the same genus: Eriocranioidea, Mimallonoidea, and Urodoidea), the average gCF is 19% and sCF 43% across the remaining superfamilies. The lowest CF values are obtained for Cossoidea (gCF=0%, sCF=33%, UFB=66) and Geometroidea (gCF=0%, sCF=35.5%, UFB=100). In the case of Cossoidea this is not so surprising because a long branch leading to *Epipomponia* (currently classified as a zygaenoid; see below for more on this group) is recovered within Cossoidea. For additional seven superfamilies, gCF values are below 5% (Bombycoidea, Drepanoidea, Gelechioidea, Papilionoidea, Pyraloidea, Tineoidea, and Zygaenoidea). We find this very surprising as all of these groups with the exception of Tineoidea and Zygaenoidea have been strongly supported as monophyletic in different studies (Kaila et al. 2011; Zwick et al. 2011; Heikkilä et al. 2012; Regier et al. 2012; Regier et al. 2013; Sohn et al. 2013; Heikkilä et al. 2014; Heikkilä et al. 2015; Espeland et al. 2018; Chazot et al. 2019).

In our closer inspection of concordance factor values in butterflies (Papilionoidea), for which the topology that we recovered is the same as the well-resolved topologies from other studies (Espeland et al. 2018; Chazot et al. 2019), we observe higher CF values compared to the backbone and on branches leading to superfamilies, but still lower than expected for such well-resolved and supported topology. In the NT12 dataset for the butterflies, we recovered the average gCF of 48.1% and sCF of 52.3% (Fig. S19), while in the NT123 the average for gCF values was higher (65.2%), but for sCF lower (46.7%) (Fig. S20). Our interpretation of this finding is that with the addition of 3^rd^ codon positions, at this shallower lever in the tree, each gene has more phylogenetic signal and that leads to the increase in the gCFs. However, the 3^rd^ codon positions, when analysed as single sites, add noise and therefore lower the sCF values.

In a comparison of the CF values from our dataset with those in published studies, we found that CF values tend to be higher in shallow phylogenies, i.e. phylogenies inferred for groups younger than 50 My, e.g. a family of frogs (Chan et al. 2020), a family of plants (Dupin et al. 2020), an ant genus (van Elst et al. 2021), and a spider family (Bond et al. 2020). The same could be observed in the results that Minh et al. (2020) obtained in the datasets that they tested for the method development. For example, in their analysis of the Misof et al. (2014) dataset for the phylogeny of insects, most gCF values are below 25% and many are close to zero. As this method is used more, it will be interesting to see whether there are datasets for deep-level phylogenies that have high concordance factors.

Minh et al. (2020) stated in the publication introducing concordance factor that it can be hard to interpret these values when they are low as two very different reasons can be responsible: 1) a strong discordance among the genes or 2) weak phylogenetic signal. Having lower gCF values than sCF values in our NT12_all dataset suggests that the lack of signal is behind the observed pattern (Lanfear 2018). We hypothesize that this stems from our gene fragments being too short (median of 705 bp) to resolve the deep branches in the lepidopteran tree-of-life. In the only other published study using CF on a phylogeny of a similar depth in time (in Aranea – spiders; Kallal et al. 2020), similar low CF values were observed overall as well as for well-established clades based on both morphological and molecular data. The same explanation that low gCF values might result from the short length of gene fragments has been offered by Minh et al. (2020) but so far there have not been any recommendations for the necessary locus length for concordance factors to be useful. This could be established for example with simulations.

In principle we agree with the strong arguments that concordance factors should be used to assess branch support in phylogenomic datasets because bootstrap values are not appropriate as they are inflated for datasets with such a high number of sites (Minh et al. 2020). However, we find that concordance factors might have limited utility for deep phylogenies when single gene fragments are relatively short, as is the case in our dataset. Reliance on concordance factors alone could undermine our confidence in branches that are otherwise very well supported by smaller molecular datasets as well as morphological synapomorpies. Therefore, we decided to fall back on exploring the robustness of our data using UFB and SH as branch support measures, as well as the average UFB (aUFB) when comparing the robustness of entire topologies.

Overall the analyses based on the nucleotide alignments resulted in higher branch supports for the backbone (Fig. 3), with the average UFB (aUFB) of 98 across all of the backbone nodes for the NT12_all dataset vs. 90 for the AA_all dataset. The best-supported AA topology was AA_nohighGC (aUFB=93; Figs 2g, S17) and the best-supported NT12 topology was NT12_no_Gele (aUFB=99; Fig. 2e, S10). With the successive removal of 10, 20, and 50 genes, as expected, aUFB somewhat decreased in the NT12 dataset (to 96, 95, and 95, respectively), but interestingly in the AA_minus10 dataset there was a small increase compared to the AA_all dataset to 91, followed by a decrease to 85 (minus20) and 87 (minus50). Removal of the 50 genes with a high GC content resulted in somewhat lower aUFB in the NT12 dataset (aUFB=94) but higher values in the AA dataset (aUFB=93) (Figs 4, S21–26). It is noteworthy that the removal of Gelechioidea improved branch support for the backbone in both NT12 (aUFB=99) and AA datasets (aUFB=92) compared to NT12_all and AA_all. NT12_degen analysis had higher backbone support than the AA but lower than the NT12 analysis (aUFB=93) (Fig. 3).

**Figure 3.**
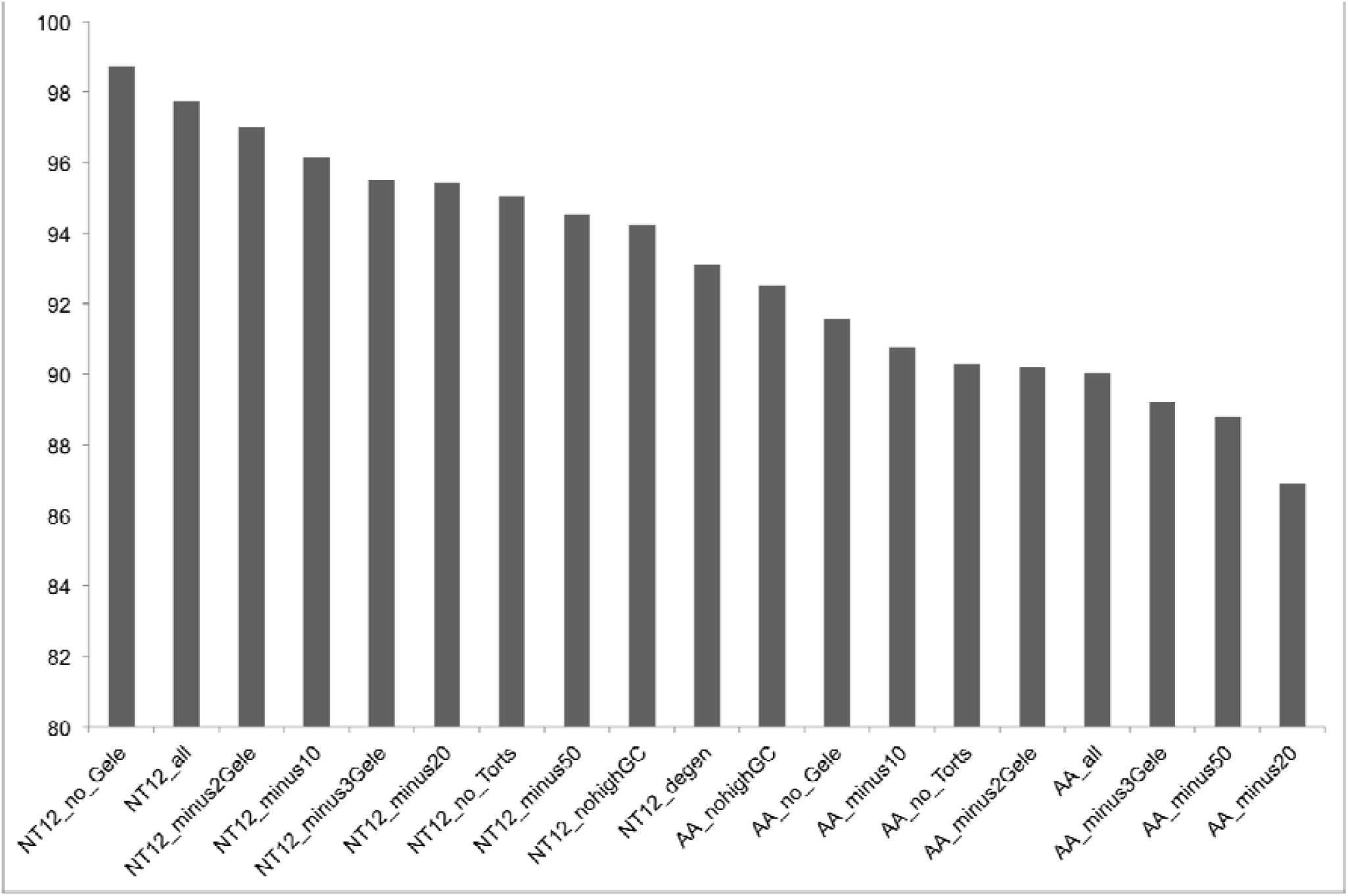
A bar graph showing the average ultrafast bootstrap (aUFB) support for the backbone nodes in trees resulting from analyses of all of the 19 datasets in this study. Note that the y-axis range is 80–100. Dataset descriptions are available in Table 1. See text for more details.

**Figure 4.**
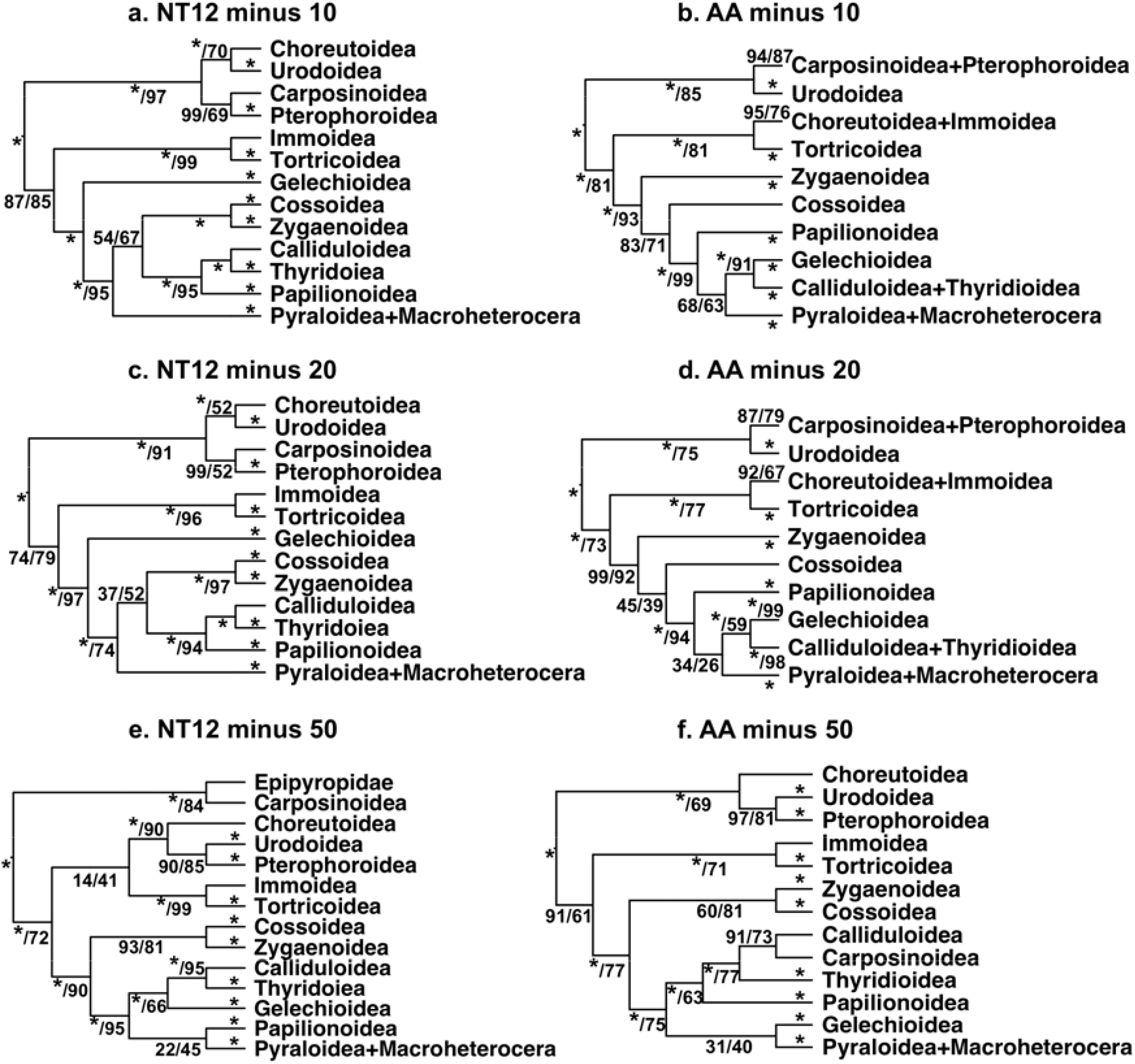
An overview of phylogenetic relationships among apoditrysian butterflies and moths from maximum likelihood analyses of different datasets. Full dataset descriptions are available in Table 1; NT12 – nucleotide alignment including 1^st^ and 2^nd^ codon positions; AA – amino acid alignment; minus10, minus20, and minus50 refer to datasets with 10, 20, and 50 genes removed, respectively, based on the strength of their phylogenetic signal. Numbers and characters on branches represent branch support as SH-like approximate likelihood ratio test / ultrafast bootstrap (UFBS), with “*” denoting 100.

A large number of clades received a UFB of 100 in all of the analyses. These are the following: Lepidoptera, Heteroneura, Eulepidoptera, Euheteroneura, Ditrysia, (Yponomeutoidea + Gracillarioidea) + Apoditrysia, and Macroheterocera (Fig. 1), as well as almost all superfamily clades. Glossata received a UFB of 100 in all except NT12_minus50 (UFB=99), while Angiospermivora received less than 100 in NT12_minus10, NT12_minus20, and NT12_minus50 (99, 97, and 99, respectively). Pyraloidea + Macroheterocera received less than 100 in AA_minus50 (99). Altogether, these clades appear to be very robust to the various perturbations of the data. Their recovery in other phylogenomic datasets (Table 3) further demonstrates the stability of these clades. We consider these parts of the lepidopteran tree of life fully resolved.

Notable exceptions to superfamilies with strongly supported monophyly are Tineoidea and Palaephatoidea. Tineoidea were recovered paraphyletic with respect to other Ditrysia in all analyses (see more details below), while Palaephatoidea are split into two clades: Palaephatoidea1 + ((Tischerioidea + Palaephatoidea2) + Ditrysia). In some of the analyses with the reduced gene sets, the branch support for some other superfamilies also decreased, e.g. Cossoidea (UFB=98 in AA_nohighGC), Neopseustoidea (UFB=97 in AA_minus10), but it remained strong across all of the analyses (UFB>95).

Removing 10 and 20 genes based on their phylogenetic signal had no effect on the NT12 topology although aUFB became somewhat lower (Fig. 4a,c). However, when 50 of these genes were removed, the topology changed in several ways with respect to the smaller apoditrysian superfamilies (Fig. 4e). For example, *Epipomponia* (Epipyropidae), which was recovered within Cossoidea in all of our other analyses and is on a long branch, was now recovered as sister to Carposinoidea (SH 100, UFB 84). Although the phylogenetic placement of Epipyropidae remains unclear, so far there have been no suggestions of a close relationship with Carposinoidea. Epipyropids are ectoparasites on homopteran insects, an uncommon lifestyle among lepidopterans. Traditionally they have been considered to belong within Zygaenoidea (Common 1970; Epstein et al. 1998) and there is possible support for this placement in morphological characters (Heikkilä et al. 2015), but in the few molecular phylogenetic studies in which epipyropids were included, they were recovered as sister to Sesiidae or Cossidae (Cossoidea) with low support and commonly on a very long branch (Mutanen et al. 2010; Bazinet et al. 2013; Regier et al. 2013). Long branches and accelerated molecular evolution have been noted in various parasitic taxa (Bernt et al. 2013). Furthermore, Epipyropidae + Carposinoidea were in turn recovered as sister to all other Apoditrysia (SH 100, UFB 72). Choreutoidea were recovered as sister (SH 100, UFB 90) to Urodoidea + Pterophoroidea (SH 90, UFB 85). The clade with these three superfamilies was recovered as sister to a clade consisting of Tortricoidea + Immoidea (SH 100, UFB 99) but without support (SH 14, UFB 41). Interestingly, Gelechioidea were recovered as sister to Calliduloidea plus Thyridoidea (SH 100, UFB 66; same position as in AA_all). The position of Papilionoidea, now recovered as sister to Pyraloidea plus Macroheterocera, but without support (SH 22, UFB 45), has not been seen in any other analysis or study. All of these anomalous placements suggest that the 50 genes with the highest phylogenetic signal are crucial for resolving the relationships among lepidopteran superfamilies.

The effect of the removal of the genes with the highest phylogenetic signal from the AA alignment was stronger. When 10 of these genes were removed, Choreutoidea changed position from being sister to Urodoidea in the AA_all dataset (SH 69.7, UFB 65) to being sister to Immoidea (SH 95, UFB 76) (Fig. 4b). With the removal of 20 genes, there were no more changes in the topology, but branch support decreased (Fig. 4d). When 50 of these genes were removed, the topology was significantly perturbed in a number of ways: Choreutoidea were recovered as sister to Urodoidea plus Pterophoroidea (SH 100, UFB 69), Zygaenoidea and Cossoidea were recovered as sisters (SH 60, UFB 81; same as in the NT12 trees), Carposinoidea were recovered as sister to Calliduloidea (SH 91, UFB 73; similar to results in Mayer et al. 2021), which were together sister to Thyridoidea (SH 100, UFB 77), all three being sister to Papilionoidea (SH 100, UFB 63), while Gelechioidea were sister to Pyraloidea plus Macroheterocera (SH 31, UFB 40) (Fig. 4f).

Furthermore, although it took more than just one gene, similar to some of the studies with contentious relationships presented by Shen et al. (2017), our AA topology also changed with removal of relatively few genes (10 genes or 3% of the total) and both the AA and NT12 topology grossly changed when 50 genes (15% of the total) were removed. As in our other analyses, the branch support overall decreased more in the trees from the AA dataset than from the NT12 dataset (Fig. 3) with removal of 10, 20, and 50 genes.

It appears that both datasets are sensitive to the removal of the genes with the highest phylogenetic content, with a stronger effect in the AA dataset, but this effect overall is not as strong as demonstrated in a number of other phylogenomic datasets by Shen et al. (2017), where removal of just a few genes resulted in the change of topology. In our dataset, 20 (in AA) or 50 genes (in NT12) need to be removed for significant changes to the topology to occur. The relationships that appear to be very strongly supported by the NT12 dataset and remain even after the removal of 50 genes are the sister relationship between Tortricoidea and Immoidea, as well as the ones between Zygaenoidea and Cossoidea, Calliduloidea and Thyridoidea, and Pyraloidea and Macroheterocera, respectively. It is worth noting that the topology from the AA_minus50 dataset is the only AA analysis of our data that includes the sister relationship between Cossoidea and Zygaenoidea, which is recovered in all of our analyses of the NT12 data, as well as in almost all other phylogenomic studies, both from AA and NT alignments, with the exception of Bazinet et al. (2013) (Table 4).

### Other sources of systematic bias

It is clear that compositional bias is behind some of the discordances in topologies resulting from nucleotide and amino acid alignments. It has been established that such bias can lead to incorrect phylogeny (e.g. Masta et al. 2009; Zwick et al. 2012). As amino acids are considered to be less susceptible to saturation in comparison to nucleotides, they have been the favoured choice of many researchers when inferring divergences that are hundreds of millions of years old (e.g. Misof et al. 2014). Using amino acids as opposed to the various RY coding schemes (such as degen1) results in more character states and a smaller information loss (Masta et al. 2009). However, if amino acid evolution is not modelled appropriately, the phylogeny inference can be misled (Gillung et al. 2018; Young and Gillung 2020) and there are cases in which unrelated groups are inferred in the same clade due to shared atypical amino acids and not their shared ancestry (Masta et al. 2009).

Studies in which amino acid coding and its impact on the phylogeny inference were explored in detail are few and far between. For example, Zwick et al. (2012) explored the effect that different ways of coding of Ser has on the topology and branch support. In the standard amino acid coding, Ser is just one state. However, in degen1 coding it is TCN and ACY. When these two sets of codons were coded as Ser1 and Ser2 in the amino acid analysis, the result was a large increase in bootstraps for the nodes of interest, demonstrating that additional signal was retrieved from this change in character coding. The issue at hand is that the transformation from Ser1 into Ser2 is a two-step process and that Ser1 and Ser2 have different rates of change into other amino acids, so synonymizing Ser1 and Ser2 leads to incorrect estimation of model parameters as well as information loss (Zwick et al. 2012). This example from Zwick et al.’s study was recounted to make the point that simply coding each amino acid without understanding their substitution processes can lead to serious problems in phylogenetic inference. Other studies have pointed out potential problems with using amino acids as characters already in the early 2000s. For example, Simmons et al. (2002) argued that amino acids are more subject to convergence as they are composite characters. It is clear that more research is needed to establish when it is appropriate to analyse amino acid alignments and when nucleotide alignments, and the reasons behind the discrepancies in results between these two types of analyses.

### Current understanding of the lepidopteran tree of life

Relationships among the paraphyletic assemblage of so-called non-ditrysian Lepidoptera appear to be settling down, with initial anomalous results being resolved with more taxa and genes being analysed (Mutanen et al. 2010; Regier et al. 2013; Kristensen et al. 2015; Regier et al. 2015b; Bazinet et al. 2017; Kawahara et al. 2019). It appears clear now that Micropterigidae are sister to the rest of Lepidoptera, followed by Agathiphagidae and then Heterobathmiidae. This classic (Kristensen 1998) arrangement was found only with phylogenomic data (Bazinet et al. 2017; Kawahara et al. 2019). Previous studies with a handful of gene sequences suggested that Agathiphagidae were sister to Heterobathmiidae (e.g. Kristensen et al. 2015; Regier et al. 2015b), a result that was strongly incongruent with morphological data. Our dataset, reduced in the number of genes but with denser taxon sampling compared to Bazinet et al. (2017), also finds this classic arrangement with strong support, as does the Kawahara et al. (2019) study.

From among the non-ditrysian superfamilies, Palaephatoidea, a group with Gondwanan distribution, stand out as being consistently recovered as two separate lineages. The resolution in this part of the tree is lacking and the important question of which group is the sister to Ditrysia (Regier et al. 2015a), the lepidopteran clade containing 98% of the order’s species diversity (van Nieukerken et al. 2011), remains unanswered. In our study the Palaephatoidea genera *Azaleodes, Metaphatus*, and *Ptyssoptera* are together sister to Tischeriidae, which are together sister to Ditrysia whereas the genus *Palaephatus* forms a separate lineage that is sister to all of these groups together. However, in other studies (Kawahara et al. 2019; Mayer et al. 2021) this relationship is reversed and it is *Palaephatus* that is recovered as the immediate sister to Ditrysia. This is yet another example of relationships where molecular evidence appears to be as confounding as the morphological (as e.g. the relationships among Neoaves in birds; Reddy et al. 2017).

The most unresolved part of the Lepidoptera tree of life is Ditrysia (Mitter et al. 2017) and especially Apoditrysia, with a number of conflicts between different studies and even different analyses of the same data. The ditrysian superfamily Tineoidea is another superfamily that requires redefinition to render the groups monophyletic. The paraphyly of Tineoidea with respect to the rest of Ditrysia has been recovered repeatedly with three separate lineages branching off sequentially, first genus *Eudarcia* (currently classified as Meessiidae), then family Psychidae, and finally the other two sampled genera, *Dryadaula* (Dryadaulidae) and *Tineola* (Tineidae) together recovered as sister to all other Ditrysia (Kawahara et al. 2019; Mayer et al. 2021; our AA_all analysis). In our NT12_all analysis the paraphyly persists because of the separate lineage leading to Meessiidae and Psychidae while Dryadaulidae + Tineidae are recovered as sister lineage to other Ditrysia. In a study with fewer genes (19 nuclear protein-coding genes) and much better taxon sampling across Tineoidea, the recovered relationships were somewhat different although Tineoidea were still paraphyletic, with Tineidae alone being sister to all other Ditrysia (Regier et al. 2015a). However, Heikkilä et al. (2015) recovered monophyletic Tineoidea when they combined the eight legacy genes (COI and seven nuclear genes; Wahlberg and Wheat 2008) with a morphological character matrix. Likely better taxon and gene sampling are necessary to put this question to rest.

Within the rest of Ditrysia, the position of Yponomeutoidea + Gracillaroidea as sister to Apoditrysia is well supported in all of the studies. There is agreement across datasets and analysis types in the position of Choreutoidea, Urodoidea, Immoidea, and Tortricoidea outside of Obtectomera, as well as the sister group relationship between Pyraloidea and Macroheterocera. Disagreement, however, persists in the recovery of the remaining apoditrysian groups. In our analyses, Pterophoroidea are never in Obtectomera, while in several other studies they are recovered within that clade (Regier et al. 2013; Kawahara et al. 2019; Mayer et al. 2021). The sister relationship of Zygaenoidea and Cossoidea is usually but not always recovered (e.g. not in our AA_all analysis). Disagreement also exists in the placement of the macroheteroceran groups with Noctuoidea being sister to (Bombycoidea + Lasiocampoidea) in our dataset analyses as well as by Heikkilä et al. (2015), while Geometroidea are recovered in the place of Noctuoidea by Kawahara et al. (2019) and Mayer et al. (2021), and even Geometroidea + Noctuoidea as sister to Bombycoidea + Lasiocampoidea by Kawahara and Breinholt (2014). Finally, as already discussed above, the position of Gelechioidea differs widely depending on the datasets and analyses carried out.

In our analyses, Gelechioidea are recovered outside of Obtectomera in the NT12 analyses except when we exclude the gelechioid taxa with a high compositional bias, which then results in the remaining gelechioids recovered within Obtectomera. In the AA analyses, Gelechoidea are recovered within Obtectomera, as sister to Calliduloidea + Thyridoidea, with this clade in turn sister to Macroheterocera. Gelechioidea are recovered within Obtectomera in several other phylogenomic studies (Kawahara et al. 2019; Mayer et al. 2021). Despite sharing two obtectomeran synapomorphies – a structure in the adult mouth parts (specifically the haustellum; Rammert 1994) as well as the pupal type, gelechioids were placed outside of that clade by Minet and others in their morphology-based but non-analytical studies (Minet 1986; Minet 1991; Kristensen and Skalski 1998). The presence of obtectomeran pupa in several other superfamilies (such as Yponomeutoidea, Epermenioidea, and Alucitoidea) was in the past interpreted as convergence (Minet 1991), but it now appears that this character is less homoplasious than previously believed. First, Heikkilä et al. (2015) refuted the presence of this character in Yponomeutoidea, and second, most of the other groups have recently been recovered within Obtectomera (Heikkilä et al. 2015; Kawahara et al. 2019; Mayer et al. 2021). However, as noted above, whether Immoidea possess an obtectomeran pupa is difficult to determine.

The position of the smaller apoditrysian superfamilies, such as Carposinoidea, Urodoidea, Choreutoidea, and Pterophoroidea, remains unresolved as their placement to some extent depends on the datasets as well as analysis type. Mitter et al. (2017) included some of these groups within Obtectomera. Of these, Pterophoroidea and Carposinoidea do not fall within Obtectomera in our analyses, but rather group with Urodoidea and Choreutoidea, well outside of Obtectomera unlike in the topologies obtained in Kawahara et al. (2019) and Mayer et al. (2021). The case of Pterophoroidea is interesting, as its position has changed wildly in previous studies. Mutanen et al. (2010) found them to be in a clade with Carposinoidea, Urodoidea, Epermenioidea among other small families outside of Obtectomera. Regier et al. (2013), on the other hand, placed them (along with Carposinoidea) as sister to Papilionoidea, well within Obtectomera. Similarly, in the phylogenomic studies, Bazinet et al. (2013), Kawahara et al. (2019), and Mayer et al. (2021) found Pterophoroidea to be within Obtectomera sister to Thyridoidea in a clade with Gelechioidea, whereas Kawahara and Breinholt (2014) found Pterophoroidea to be sister to Urodoidea in a clade with Tortricoidea. Clearly, a more careful investigation is needed to establish where this group belongs.

Likewise, the position of another small superfamily, Choreutoidea, has been elusive for a long time. Since their description, Choreutoidea had been placed in four different superfamilies (review in Rota 2011) before they were placed in their own monotypic superfamily Choreutoidea (Minet 1991). Morphologically they are considered to belong to Apoditrysia (Kristensen 1998). In the two major phylogenetic studies they grouped with completely different families (with Immoidea, Galacticidae, and Epipyropidae in Mutanen et al. (2010); with Douglasiidae and Schreckensteiniidae in Regier et al. (2013)) without support. In our study as well as the other phylogenomic studies they tend to be recovered in a clade with Urodoidea but the membership of this clade changes widely between different studies and analysis type. Several other smaller lepidopteran superfamilies, such as Galacticoidea, Hyblaeoidea, Schreckensteinioidea Simaethistoidea, as well as unplaced families Millieriidae and Prodidactidae have not been sampled in any of the phylogenomic studies and for some of them we have no molecular data at all. Including these groups in future phylogenomic studies could potentially shed more light on this part of the lepidopteran tree of life and that should be among priorities for lepidopteran systematists.

### Concluding remarks

“The real test of whether phylogenomics can fulfil the promise to resolve the tree of life will depend on careful scrutiny of the data for patterns of sequence evolution that might lead to bias and understanding the impact of those patterns on the results of phylogenetic analyses.” (Reddy et al. 2017)

Despite significant progress in understanding the phylogenetic relationships of the major lineages of the megadiverse order Lepidoptera in the last decade, our analyses show that there are still major issues that need to be resolved. In particular, it is now clear that the backbone of Ditrysia is still largely unresolved. Interestingly, the stable parts of Ditrysia were already picked up by Sanger sequencing studies (Regier et al. 2009; Mutanen et al. 2010; Regier et al. 2013), and despite article titles to the contrary (Kawahara and Breinholt 2014; Kawahara et al. 2019), our knowledge of the relationships of the remaining parts of Ditrysia is still murky.

It is especially disconcerting that the overall topology of Ditrysia is very different depending on the dataset as well as to some extent on whether the analyses are carried out on the AA or NT12 alignments and that the different placement of some groups, e.g. the large microlepidopteran superfamily Gelechioidea, is strongly supported in both sets of analyses. However, there is no clear answer on whether analysing amino acid or nucleotide alignments is better for phylogeny inference at this level (i.e. insect order). In seven out of 13 phylogenomic studies of insect orders the analyses were done on both kinds of data and in five of these the results were conflicting (literature review; Table 2). Often the authors choose to present results from only one kind of data without necessarily providing rationale for their choice even when they note that there is conflict.

Some studies have demonstrated that obtaining real phylogenetic signal from amino acid alignments is not simple and requires recoding of the amino acids (e.g., Masta et al. 2009; Zwick et al. 2012). The problem is that it is not *a priori* clear how to recode the data as that depends on the nucleotide bias as well as amino acid bias specific to the dataset at hand. Another problem is that the available phylogenetic inference programs cannot accommodate such recoded datasets: e.g. coding serine as two different states depending on whether the codons for it are TCN or ACY as was done by Zwick et al. (2012).

Instead of studying the amino acid composition and potential biases in our data, we focused on establishing whether the results from amino acid or nucleotide data are more robust. Our findings clearly demonstrate that the nucleotide analyses are more robust if robustness is measured with nodal support and topological stability. The overall measure of backbone nodal support that we employed – aUFB – was always higher in the NT12 than in the AA datasets (Fig. 3). Removing Gelechioidea from the NT12 topology did not change the topology with respect to all other clades whereas the resulting AA topology without Gelechioidea actually changed so that it became more similar to the NT12_all topology with respect to the position of Calliduloidea and Thyridoidea. Likewise, removal of the 50 genes with high GC content resulted in the same topology for the NT12 dataset but a different one for the AA dataset, while topologies from both datasets changed in different ways with the removal of Tortricoidea.

It appears that Gelechioidea are driving the main differences in the topology between the amino acid and nucleotide analyses. It is likely that the high compositional bias of gelechioid taxa in both AA and NT12 datasets is responsible for the different positions of this clade. If this is the main reason, then the degen1 analysis might be the best hypothesis at this point since degen1 coding is supposed to be relatively insensitive to the composition bias (Zwick et al. 2012). Another possible explanation comes from the large undersampling of this superfamily – we have representatives of only four of the 21 gelechioid families and other Lepidoptera phylogenomic studies are based on a similarly low sampling.

Although Lepidoptera are relatively well sampled compared to other insect groups, and even most animal groups, we believe that what is needed at this stage to improve the resolution of the tree of life of butterflies and moths is an increase in taxon sampling even if it means sampling only hundreds of genes instead of thousands. Recent studies have given us tool sets for doing just that by using specimens in museum collections (Twort et al. 2020; Call et al. 2021; Mayer et al. 2021) and we expect that museomics will play an increasingly important role in phylogenomics.

We fully agree with Young and Gillung’s (2020) statement that “the future of phylogenomic analysis is extremely bright, with many independent avenues of exploration available.” In addition to seeing more studies including careful exploration of data, we also hope to see simulations or other kinds of studies that can fully answer the question of why analyses of nucleotides and amino acids tend to differ.

## Supporting information

Table S1. Taxon list.

Table S2. List of genes analysed in this study with various statistics on them.

Table S3. Literature review.

## Acknowledgements

We are grateful to Maria Heikkilä for reading the manuscript and providing valuable insight. AC was funded by a Crafoord Foundation (Sweden) grant awarded to JR (grant no. 20180794). NW acknowledges funding from the Swedish Research Council (grant no. 2015-0444).

## List of supplementary files

**Supplementary file 1**. Table listing information on the literature review we carried out with full references listed below the table.

Table S1. Taxon list.

Table S2. List of genes analysed in this study with various statistics on them. Table S3. Literature review.

Supplementary figures S1–S26.

## References

Ababneh F., Jermiin L.S., Ma C., Robinson J. 2006. Matched-pairs tests of homogeneity with applications to homologous nucleotide sequences. Bioinformatics, 22:1225–1231. doi: 10.1093/bioinformatics/btl064.

Abascal F., Zardoya R., Telford M.J. 2010. TranslatorX: multiple alignment of nucleotide sequences guided by amino acid translations. Nucleic Acids Research, 38:W7–13. doi: 10.1093/nar/gkq291.

Altschul S.F., Gish W., Miller W., Myers E.W., Lipman D.J. 1990. Basic local alignment search tool. Journal of Molecular Biology, 215:403-410. doi.

Bazinet A.L., Cummings M.P., Mitter K.T., Mitter C.W. 2013. Can RNA-Seq resolve the rapid radiation of advanced moths and butterflies (Hexapoda: Lepidoptera: Apoditrysia)? An exploratory study. PLoS ONE, 8:e82615. doi: 10.1371/journal.pone.0082615.

Bazinet A.L., Mitter K.T., Davis D.R., van Nieukerken E.J., Cummings M.P., Mitter C. 2017. Phylotranscriptomics resolves ancient divergences in the Lepidoptera. Systematic Entomology, 42:305–316. doi: 10.1111/syen.12217.

Bernt M., Bleidorn C., Braband A., Dambach J., Donath A., Fritzsch G., Golombek A., Hadrys H., Jühling F., Meusemann K., et al. 2013. A comprehensive analysis of bilaterian mitochondrial genomes and phylogeny. Molecular Phylogenetics and Evolution, 69:352–364. doi: 10.1016/j.ympev.2013.05.002.

Bond J.E., Hamilton C.A., Godwin R.L., Ledford J.M., Starrett J. 2020. Phylogeny, evolution, and biogeography of the North American trapdoor spider family Euctenizidae (Araneae: Mygalomorphae) and the discovery of a new ‘endangered living fossil’ along California’s central coast. Insect Systematics and Diversity, 4:5:2. doi: 10.1093/isd/ixaa010.

Bowker A.H. 1948. A test for symmetry in contingency tables. Journal of the American Statistical Association, 43:572–574. doi: 10.1080/01621459.1948.10483284.

Breinholt J.W., Earl C., Lemmon A.R., Lemmon E.M., Xiao L., Kawahara A.Y. 2018. Resolving relationships among the megadiverse butterflies and moths with a novel pipeline for anchored phylogenomics. Systematic Biology, 67:78–93. doi: 10.1093/sysbio/syx048.

Call E., Mayer C., Dietz L., Twort V.G., Wahlberg N., Espeland M. 2021. Museomics: phylogenomics of the moth family Epicopeiidae (Lepidoptera) using target enrichment. Insect Systematics & Diversity, in press. doi.

Chan K.O., Hutter C.R., Wood P.L., Grismer L.L., Brown R.M. 2020. Target-capture phylogenomics provide insights on gene and species tree discordances in Old World treefrogs (Anura: Rhacophoridae). Proceedings of the Royal Society B: Biological Sciences, 287:20202102. doi: 10.1098/rspb.2020.2102.

Chazot N., Wahlberg N., Freitas A.V.L., Mitter C., Labandeira C., Sohn J.-C., Sahoo R.K., Seraphim N., de Jong R., Heikkilä M. 2019. Priors and posteriors in Bayesian timing of divergence analyses: the age of butterflies revisited. Systematic Biology, 68:797–813. doi: 10.1093/sysbio/syz002.

Common I.F.B. 1970. Lepidoptera (Moths and Butterflies). In: CSIRO editor. The Insects of Australia. Melbourne, Australia, Melbourne Univ. Press, p. 765–866.

Di Franco A., Poujol R., Baurain D., Philippe H. 2019. Evaluating the usefulness of alignment filtering methods to reduce the impact of errors on evolutionary inferences. BMC Evolutionary Biology, 19:21. doi: 10.1186/s12862-019-1350-2.

Dupin J., Raimondeau P., Hong-Wa C., Manzi S., Gaudeul M., Besnard G. 2020. Resolving the phylogeny of the olive family (Oleaceae): confronting information from organellar and nuclear genomes. Genes, 11. doi: 10.3390/genes11121508.

Epstein M.E., Geertsema H., Naumann C.M., Tarmann G.E. 1998. The Zygaenoidea. In: Kristensen NP editor. Lepidoptera, Moths and Butterflies, 1: Evolution, Systematics, and Biogeography.. Berlin, Walter de Gruyter, p. 169–170.

Espeland M., Breinholt J., Willmott K.R., Warren A.D., Vila R., Toussaint E.F.A., Maunsell S.C., Aduse-Poku K., Talavera G., Eastwood R., et al. 2018. A comprehensive and dated phylogenomic analysis of butterflies. Current Biology, 28:P770–P778. doi: 10.1016/j.cub.2018.01.061.

Gillung J.P., Winterton S.L., Bayless K.M., Khouri Z., Borowiec M.L., Yeates D., Kimsey L.S., Misof B., Shin S., Zhou X., et al. 2018. Anchored phylogenomics unravels the evolution of spider flies (Diptera, Acroceridae) and reveals discordance between nucleotides and amino acids. Molecular Phylogenetics and Evolution, 128:233–245. doi: 10.1016/j.ympev.2018.08.007.

Grabherr M.G., Haas B.J., Yassour M., Levin J.Z., Thompson D.A., Amit I., Adiconis X., Fan L., Raychowdhury R., Zeng Q.D., et al. 2011. Full-length transcriptome assembly from RNA-Seq data without a reference genome. Nature Biotechnology, 29:644–U130. doi: 10.1038/nbt.1883.

Guindon S., Dufayard J.-F., Lefort V., Anisimova M., Hordijk W., Gascuel O. 2010. New algorithms and methods to estimate Maximum-Likelihood phylogenies: Assessing the performance of PhyML 3.0. Systematic Biology, 59:307–321. doi: 10.1093/sysbio/syq010.

Haas B.J., Papanicolauo A., Yassour M., Grabherr M.G., Blood P.D., Bowden J., Cougar M.B., Eccles D., Li B., Lieber M., et al. 2013. De novo transcript sequence reconstruction from RNA-seq using the Trinity platform for reference generation and analysis. Nature Protocols, 8:1494–1512. doi: 10.1038/nprot.2013.084.

Heikkilä M., Kaila L., Mutanen M., Peña C., Wahlberg N. 2012. Cretaceous origin and repeated Tertiary diversification of the redefined butterflies. Proceedings of the Royal Society of London B Biological Sciences, 279:1093–1099. doi: 10.1098/rspb.2011.1430.

Heikkilä M., Mutanen M., Kekkonen M., Kaila L. 2014. Morphology reinforces proposed molecular phylogenetic affinities: a revised classification for Gelechioidea (Lepidoptera). Cladistics, 30:563–589. doi: 10.1111/cla.12064.

Heikkilä M., Mutanen M., Wahlberg N., Sihvonen P., Kaila L. 2015. Elusive ditrysian phylogeny: an account of combining systematized morphology with molecular data (Lepidoptera). BMC Evolutionary Biology, 15:260. doi: 10.1186/s12862-015-0520-0.

Hoang D.T., Chernomor O., von Haeseler A., Minh B.Q., Vinh L.S. 2018. UFBoot2: Improving the ultrafast bootstrap approximation. Molecular Biology and Evolution, 35:518–522. doi: 10.1093/molbev/msx281.

Jermiin L.S., Ho S.Y.W., Ababneh F., Robinson J., Larkum A.W.D. 2004. The biasing effect of compositional heterogeneity on phylogenetic estimates may be underestimated. Systematic Biology, 53:638–643. doi: 10.1080/10635150490468648.

Kaila L. 2004. Phylogeny of the superfamily Gelechioidea (Lepidoptera: Ditrysia): an examplar approach. Cladistics, 20:303-340. doi.

Kaila L., Mutanen M., Nyman T. 2011. Phylogeny of the mega-diverse Gelechioidea (Lepidoptera): Adaptations and determinants of success. Molecular Phylogenetics and Evolution, 61:801–809. doi: 10.1016/j.ympev.2011.08.016.

Kaila L., Nupponen K., Gorbunov P.Y., Mutanen M., Heikkilä M. 2020. Ustyurtiidae, a new family of Urodoidea with description of a new genus and two species from Kazakhstan, and discussion on possible affinity of Urodoidea to Schreckensteinioidea (Lepidoptera). Insect Systematics & Evolution, 51:444–471. doi: 10.1163/1876312X-00002209.

Kallal R.J., Kulkarni S.S., Dimitrov D., Benavides L.R., Arnedo M.A., Giribet G., Hormiga G. 2020. Converging on the orb: denser taxon sampling elucidates spider phylogeny and new analytical methods support repeated evolution of the orb web. Cladistics, in press. doi: 10.1111/cla.12439.

Kalyaanamoorthy S., Minh B.Q., Wong T.K.F., von Haeseler A., Jermiin L.S. 2017. ModelFinder: Fast model selection for accurate phylogenetic estimates. Nature Methods, 14:587–589. doi: 10.1038/nmeth.4285.

Katoh K., Kuma K.-i., Toh H., Miyata T. 2005. MAFFT version 5: improvement in accuracy of multiple sequence alignment. Nucleic Acids Research, 33:511–518. doi: 10.1093/nar/gki198.

Katoh K., Standley D.M. 2013. MAFFT Multiple sequence alignment software version 7: improvements in performance and usability. Molecular Biology and Evolution, 30:772–780. doi: 10.1093/molbev/mst010.

Kawahara A.Y., Breinholt J.W. 2014. Phylogenomics provides strong evidence for relationships of butterflies and moths. Proceedings of the Royal Society of London B Biological Sciences, 281:20140970. doi: 10.1098/rspb.2014.0970.

Kawahara A.Y., Plotkin D., Espeland M., Meusemann K., Toussaint E.F.A., Donath A., Gimnich F., Frandsen P.B., Zwick A., dos Reis M., et al. 2019. Phylogenomics reveals the evolutionary timing and pattern of butterflies and moths. Proceedings of the National Academy of Sciences of the United States of America, 116:22657–22663. doi: 10.1073/pnas.1907847116.

Kitts P.A., Church D.M., Thibaud-Nissen F., Choi J., Hem V., Sapojnikov V., Smith R.G., Tatusova T., Xiang C., Zherikov A., et al. 2016. Assembly: a resource for assembled genomes at NCBI. Nucleic Acids Research, 44:D73–D80. doi: 10.1093/nar/gkv1226.

Kristensen N.P. 1998. Lepidoptera, Moths and Butterflies. 1. Evolution, Systematics and Biogeography. Handbook of Zoology 4 (35), Lepidoptera. Berlin, de Gruyter.

Kristensen N.P., Hilton D.J., Kallies A., Milla L., Rota J., Wahlberg N., Wilcox S.A., Glatz R.V., Young D.A., Cocking G., et al. 2015. A new extant family of primitive moths from Kangaroo Island, Australia and its significance for understanding early Lepidoptera evolution. Systematic Entomology, 40:5–16. doi: 10.1111/syen.12115.

Kristensen N.P., Skalski A.W. 1998. Phylogeny and palaeontology. In: Kristensen NP editor. Handbook of zoology, vol. IV, Arthropoda: Insecta, part 35, Lepidoptera, moths and butterflies, vol. 1.Berlin, Germany, Walter de Gruyter.

Lanfear R. 2018. Calculating and interpreting gene-and site-concordance factors in phylogenomics.

Leinonen R., Sugawara H., Shumway M. 2011. The Sequence Read Archive. Nucleic Acids Research, 39:D19–D21. doi: 10.1093/nar/gkq1019.

Li W., Godzik A. 2006. Cd-hit: a fast program for clustering and comparing large sets of protein or nucleotide sequences. Bioinformatics, 22:1658–1659. doi: 10.1093/bioinformatics/btl158.

Martin M. 2011. Cutadapt removes adapter sequences from high-throughput sequencing reads. EMBnet.Journal, 17:10–11. doi: 10.14806/ej.17.1.200.

Masta S.E., Longhorn S.J., Boore J.L. 2009. Arachnid relationships based on mitochondrial genomes: Asymmetric nucleotide and amino acid bias affects phylogenetic analyses. Molecular Phylogenetics and Evolution, 50:117–128. doi: 10.1016/j.ympev.2008.10.010.

Mayer C., Dietz L., Call E., Kukowka S., Martin S., Espeland M. 2021. Adding leaves to the Lepidoptera tree: capturing hundreds of nuclear genes from old museum specimens. Systematic Entomology, in press. doi: 10.1111/syen.12481.

Miller M.A., Pfeiffer W., Schwartz T. 2010. Creating the CIPRES Science Gateway for inference of large phylogenetic trees. Proceedings of the Gateway Computing Environments Workshop (GCE). New Orleans, Louisiana.

Minet J. 1986. Ebauche d’une classification moderne de l’ordre des lépidoptères. Alexanor, 14:291-313. doi.

Minet J. 1991. Tentative reconstruction the ditrysian phylogeny (Lepidoptera: Glossata). Entomologica Scandinavica, 22:69–95. doi: 10.1163/187631291X00327.

Minh B.Q., Hahn M.W., Lanfear R. 2020. New methods to calculate concordance factors for phylogenomic datasets. Molecular Biology and Evolution, 37:2727–2733. doi: 10.1093/molbev/msaa106.

Misof B., Liu S.L., Meusemann K., Peters R.S., Donath A., Mayer C., Frandsen P.B., Ware J., Flouri T., Beutel R.G., et al. 2014. Phylogenomics resolves the timing and pattern of insect evolution. Science, 346:763–767. doi: 10.1126/science.1257570.

Mitter C., Davis D.R., Cummings M.P. 2017. Phylogeny and evolution of Lepidoptera. Annual Review of Entomology, 62:265–283. doi: 10.1146/annurev-ento-031616-035125.

Mutanen M., Wahlberg N., Kaila L. 2010. Comprehensive gene and taxon coverage elucidates radiation patterns in moths and butterflies. Proceedings of the Royal Society of London B Biological Sciences, 277:2839–2848. doi: 10.1098/rspb.2010.0392.

Nguyen L.-T., Schmidt H.A., von Haeseler A., Minh B.Q. 2015. IQ-TREE: A fast and effective stochastic algorithm for estimating maximum likelihood phylogenies. Molecular Biology and Evolution, 32:268–274. doi: 10.1093/molbev/msu300.

Nieukerken E.J.v., Kaila L., Kitching I.J., Kristensen N.P., Lees D.C., Minet J., Mitter C., Mutanen M., Regier J.C., Simonsen T.J., et al. 2011. Order Lepidoptera. In: Zhang Z-Qeditor. Animal biodiversity: An outline of higher-level classification and survey of taxonomic richness, Zootaxa, p. 212–221.

Peña C., Malm T. 2012. VoSeq: a Voucher and DNA Sequence Web Application. PlosOne, 7:e39071. doi: 10.1371/journal.pone.0039071.

Philippe H., Brinkmann H., Lavrov D.V., Littlewood D.T.J., Manuel M., Wörheide G., Baurain D. 2011. Resolving difficult phylogenetic questions: why more sequences are not enough. PLoS Biology, 9:e1000602. doi: 10.1371/journal.pbio.1000602.

Portik D.M., Smith L.L., Bi K. 2016. An evaluation of transcriptome‐based exon capture for frog phylogenomics across multiple scales of divergence (Class: Amphibia, Order: Anura). Molecular Ecology Resources, 16:1069–1083. doi: 10.1111/1755-0998.12541.

Rajaei H.S., Greve C., Letsch H., Stüning D., Wahlberg N., Minet J., Misof B. 2015. Advances in Geometroidea phylogeny, with characterization of a new family based on Pseudobiston pinratanai (Lepidoptera, Glossata). Zoologica Scripta, 44:418–436. doi: 10.1111/zsc.12108.

Rammert U. 1994. Morphologische Untersuchungen zur Aufdeckung der stammesgeschichtlichen Verhältnisse der basalen Gruppen der ditrysen Lepidopteren (Lepidoptera: Ditrysia). Universität Bielefeld, Flintbeck, Germany, p. 193.

Reddy S., Kimball R.T., Pandey A., Hosner P.A., Braun M.J., Hackett S.J., Han K.-L., Harshman J., Huddleston C.J., Kingston S., et al. 2017. Why do phylogenomic data sets yield conflicting trees? Data type influences the avian tree of life more than taxon sampling. Systematic Biology, 66:857–879. doi: 10.1093/sysbio/syx041.

Regier J.C., Mitter C., Davis D.R., Harrison T.L., Sohn J.-C., Cummings M.P., Zwick A., Mitter K.T. 2015a. A molecular phylogeny and revised classification for the oldest ditrysian moth lineages (Lepidoptera: Tineoidea), with implications for ancestral feeding habits of the mega-diverse Ditrysia. Systematic Entomology, 40:409–432. doi: 10.1111/syen.12110.

Regier J.C., Mitter C., Kristensen N.P., Davis D.R., van Nieukerken E.J., Rota J., Simonsen T.J., Mitter K.T., Kawahara A.Y., Yen S.-H., et al. 2015b. A molecular phylogeny for the oldest (non-ditrysian) lineages of extant Lepidoptera, with implications for classification, comparative morphology and life history evolution. Systematic Entomology, 40:671–704. doi: 10.1111/syen.12129.

Regier J.C., Mitter C., Solis M.A., Hayden J.E., Landry B., Nuss M., Simonsen T.J., Yen S.H., Zwick A., Cummings M.P. 2012. A molecular phylogeny for the pyraloid moths (Lepidoptera: Pyraloidea) and its implications for higher-level classification. Systematic Entomology, 37:635–656. doi: 10.1111/j.1365-3113.2012.00641.x.

Regier J.C., Mitter C., Zwick A., Bazinet A.L., Cummings M.P., Kawahara A.Y., Sohn J.-C., Zwickl D.J., Cho S., Davis D.R., et al. 2013. A large-scale, higher-level, molecular phylogenetic study of the insect order Lepidoptera (moths and butterflies). PlosOne, 8:e58568. doi: 10.1371/journal.pone.0058568.

Regier J.C., Shultz J.W., Zwick A., Hussey A., Ball B., Wetzer R., Martin J.W., Cunningham C.W. 2010. Arthropod relationships revealed by phylogenomic analysis of nuclear protein-coding sequences. Nature, 463:1079–1083. doi: 10.1038/nature08742.

Regier J.C., Zwick A., Cummings M.P., Kawahara A.Y., Cho S., Weller S.J., Roe A.D., Baixeras J., Brown J.W., Parr C.S., et al. 2009. Toward reconstructing the evolution of advanced moths and butterflies (Lepidoptera: Ditrysia): an initial molecular study. BMC Evolutionary Biology, 9:280. doi: 10.1186/1471-2148-9-280.

Rota J. 2011. Data partitioning in Bayesian analysis: molecular phylogenetics of metalmark moths (Lepidoptera: Choreutidae). Systematic Entomology, 36:317–329. doi: 10.1111/j.1365-3113.2010.00563.x.

Schmieder R., Edwards R. 2011. Quality control and preprocessing of metagenomic datasets. Bioinformatics. Bioinformatics, 27:863–864. doi: 10.1093/bioinformatics/btr026.

Shen X.-X., Hittinger C.T., Rokas A. 2017. Contentious relationships in phylogenomic studies can be driven by a handful of genes. Nature Ecology & Evolution, 1:0126. doi: 10.1038/s41559-017-0126.

Simmons M.P., Ochoterena H., Freudenstein J.V. 2002. Conflict between amino acid and nucleotide characters. Cladistics, 18:200–206. doi: 10.1111/j.1096-0031.2002.tb00148.x.

Sohn J.C., Regier J.C., Mitter C., Davis D., Landry J.-F., Zwick A., Cummings M.P. 2013. A molecular phylogeny for Yponomeutoidea (Insecta, Lepidoptera, Ditrysia) and its implications for classification, biogeography and the evolution of host plant use. PlosOne, 8:e55066. doi: 10.1371/journal.pone.0055066.

Stamatakis A. 2014. RAxML Version 8: A tool for phylogenetic analysis and post-analysis of large phylogenies. Bioinformatics, 30:1312–1313. doi: 10.1093/bioinformatics/btu033.

Suyama M., Torrents D., Bork P. 2006. PAL2NAL: robust conversion of protein sequence alignments into the corresponding codon alignments. Nucleic Acids Research, 34:W609–W612. doi: 10.1093/nar/gkl315.

Twort V.G., Minet J., Wheat C.W., Wahlberg N. 2020. Museomics of a rare taxon: placing Whalleyanidae in the Lepidoptera Tree of Life. BioRxiv. doi: 10.1101/2020.08.18.255182.

van Elst T., Eriksson T.H., Gadau J., Johnson R.A., Rabeling C., Taylor J.E., Borowiec M.L. 2021. Comprehensive phylogeny of Myrmecocystus honey ants highlights cryptic diversity and infers evolution during aridification of the American Southwest. Molecular Phylogenetics and Evolution, 155:107036. doi: 10.1016/j.ympev.2020.107036.

Vasilikopoulos A., Balke M., Beutel R.G., Donath A., Podsiadlowski L., Pflug J.M., Waterhouse R.M., Meusemann K., Peters R.S., Escalona H.E., et al. 2019. Phylogenomics of the superfamily Dytiscoidea (Coleoptera: Adephaga) with an evaluation of phylogenetic conflict and systematic error. Molecular Phylogenetics and Evolution, 135:270–285. doi: 10.1016/j.ympev.2019.02.022.

Wahlberg N., Wheat C.W. 2008. Genomic outposts serve the phylogenomic pioneers: designing novel nuclear markers for genomic DNA extractions of Lepidoptera. Systematic Biology, 57:231–242. doi: 10.1080/10635150802033006.

Wahlberg N., Wheat C.W., Peña C. 2013. Timing and patterns in the taxonomic diversification of Lepidoptera (butterflies and moths). PLoS ONE, 8:e80875. doi: 10.1371/journal.pone.0080875.

Wong K.M., Suchard M.A., Huelsenbeck J.P. 2008. Alignment uncertainty and genomic analysis. Science, 319:473-476. doi.

Young A.D., Gillung J.P. 2020. Phylogenomics — principles, opportunities and pitfalls of big‐data phylogenetics. Systematic Entomology, 45:225–247. doi: 10.1111/syen.12406.

Zhang C., Rabiee M., Sayyari E., Mirarab S. 2018. ASTRAL-III: polynomial time species tree reconstruction from partially resolved gene trees. BMC Bioinformatics, 19:153. doi: 10.1186/s12859-018-2129-y.

Zwick A., Regier J.C., Mitter C., Cummings M.P. 2011. Increased gene sampling yields robust support for higher-level clades within Bombycoidea (Lepidoptera). Systematic Entomology, 36:31–43. doi: 10.1111/j.1365-3113.2010.00543.x.

Zwick A., Regier J.C., Zwickl D.J. 2012. Resolving discrepancy between nucleotides and amino acids in deep-level arthropod phylogenomics: differentiating serine codons in 21-amino-acid models. PLoS ONE, 7:e47450. doi: 10.1371/journal.pone.0047450.

