## Supplementary file 1 for "The unresolved phylogenomic tree of butterflies and moths (Lepidoptera): assessing the potential causes and consequences"

**Supplementary file 1.** Table listing information on the literature review we carried out with full references listed below the table. AA refers to amino acid alignments, NT refers to nucleotide alignments, with numbers referring to codon positions, degen refers to special coding of nucleotides that only retains information from non-synonymous changes.

| **Study (author, year)** | **Journal** | **Insect group** | **Taxonomic level** | **Analysis** | **Results shown** | **Conflict present** |
| --- | --- | --- | --- | --- | --- | --- |
| Evangelista et al. 2019 | Proc Royal Soc B | Blattodea | order | AA, NT2 | AA | yes |
| Winterton et al. 2019 | Systematic Entomology | Chrysopidae | family | NT123 |  |  |
| McKenna et al. 2020 | PNAS | Coleoptera | order | AA | AA |  |
| Gustafson et al. 2020 | Systematic Entomology | Coleoptera: Adephaga | suborder | NT (UCE) |  |  |
| Cai et al. 2020 | MPE | Coleoptera: Dytiscoidea | superfamily | AA, NT123 | AA | yes |
| Vasilikopoulos et al. 2019 | MPE | Coleoptera: Dytiscoidea | superfamily | AA, NT123, NT12, degen |  | yes |
| Kusy et al. 2019 | Systematic Entomology | Coleoptera: Lycidae | family | AA, NT123 | both | yes |
| Wipfler et al. 2020 | Systematic Entomology | Dermaptera | order | AA |  |  |
| Kutty et al. 2019 | Cladistics | Diptera: Calyptratae | suborder | AA, NT123, NT2 | AA | yes |
| Buenaventura et al. 2020 | Systematic Entomology | Diptera: Sarcophagidae | family | NT123 |  |  |
| Borowiec 2019 | Systematic Biology | Formicidae | subfamily | AA, NT123 | both | yes |
| Johnson et al. 2018 | PNAS | Hemiptera | order | AA, NT12 | NT12 | no |
| Kieran et al. 2019 | MPE | Hemiptera | order | NT (UCE incl. Prot-cod) |  |  |
| Skinner et al. 2020 | Systematic Entomology | Hemiptera: Auchenorrhyncha | suborder | AA, NT123, degen | AA, NT123 | yes |
| Knyshov et al. 2019 | Systematic Entomology | Hemiptera: Schizopteridae | family | NT123 |  |  |
| Bossert et al. 2019 | MPE | Hymenoptera: Apidae | family | NT (UCE incl. Prot-cod) |  |  |
| Baker et al. 2020 | Systematic Entomology | Hymenoptera: Eucharitidae: Oraseminae | subfamily | NT123 |  |  |
| Kawahara & Breinholt 2014 | Proc Royal Soc B | Lepidoptera | order | AA, NT123 | NT123 | no |
| Kawahara et al. 2019 | PNAS | Lepidoptera | order | AA, NT123 degen | AA | no |
| Breinholt et al. 2018 | Systematic Biology | Lepidoptera | order | AA, NT12 degen | AA, NT12 degen | yes |
| Bazinet et al. 2013 | PLOS One | Lepidoptera | order | NT123 degen | NT123 degen |  |
| Bazinet et al. 2017 | Systematic Entomology | Lepidoptera | order | NT123, NT123 degen | NT123, NT123 degen |  |
| Hamilton et al. 2019 | BMC Evolutionary Biology | Lepidoptera: Bombycoidea | superfamily | AA, NT123 | NT123 | no |
| Homziak et al. 2019 | MPE | Lepidoptera: Erebinae | subfamily | NT123 |  |  |
| Zhang et al. 2020 | MPE | Lepidoptera: Geometroidea: Epicopeidae | family | AA, NT123, degen | NT123 | no |
| Milla et al. 2020 | Systematic Entomology | Lepidoptera: Heliozelidae | family | NT123 |  |  |
| St Laurent et al. 2020 | Systematic Entomology | Lepidoptera: Mimallonidae | family | NT123 |  |  |
| Espeland et al. 2019 | MPE | Lepidoptera: Nymphalidae: Satyrinae: Euptychina | tribe | NT123 |  |  |
| Machado et al. 2019 | Systematic Entomology | Myrmeleontidae | family | NT123 |  |  |
| Vasilikopoulos et al. 2019 | BMC Evolutionary Biology | Neuropterida | order | AA | AA |  |
| Allio et al. 2020 | Systematic Biology | Papilionidae | family | AA, NT123 | both | No |
| Chazot et al. 2019 | Systematic Biology | Papilionoidea | superfamily | NT123 |  |  |
| Simon et al. 2019 | Frontiers in Ecol Evol | Phasmatodea | order | AA, NT2 | AA | yes |
| Wipfler et al. 2019 | PNAS | Polyneoptera | order | AA, NT2 | AA | yes |

**References:**
