## Supplementary figures S1-S26 for "The unresolved phylogenomic tree of butterflies and moths (Lepidoptera): assessing the potential causes and consequences"

**Note:** Branch support is shown on branches with the SH-like approximate likelihood ratio test value as first number and the ultrafast bootstrap value as the second number (Sh-like/UFB) unless otherwise noted in the figure legend.

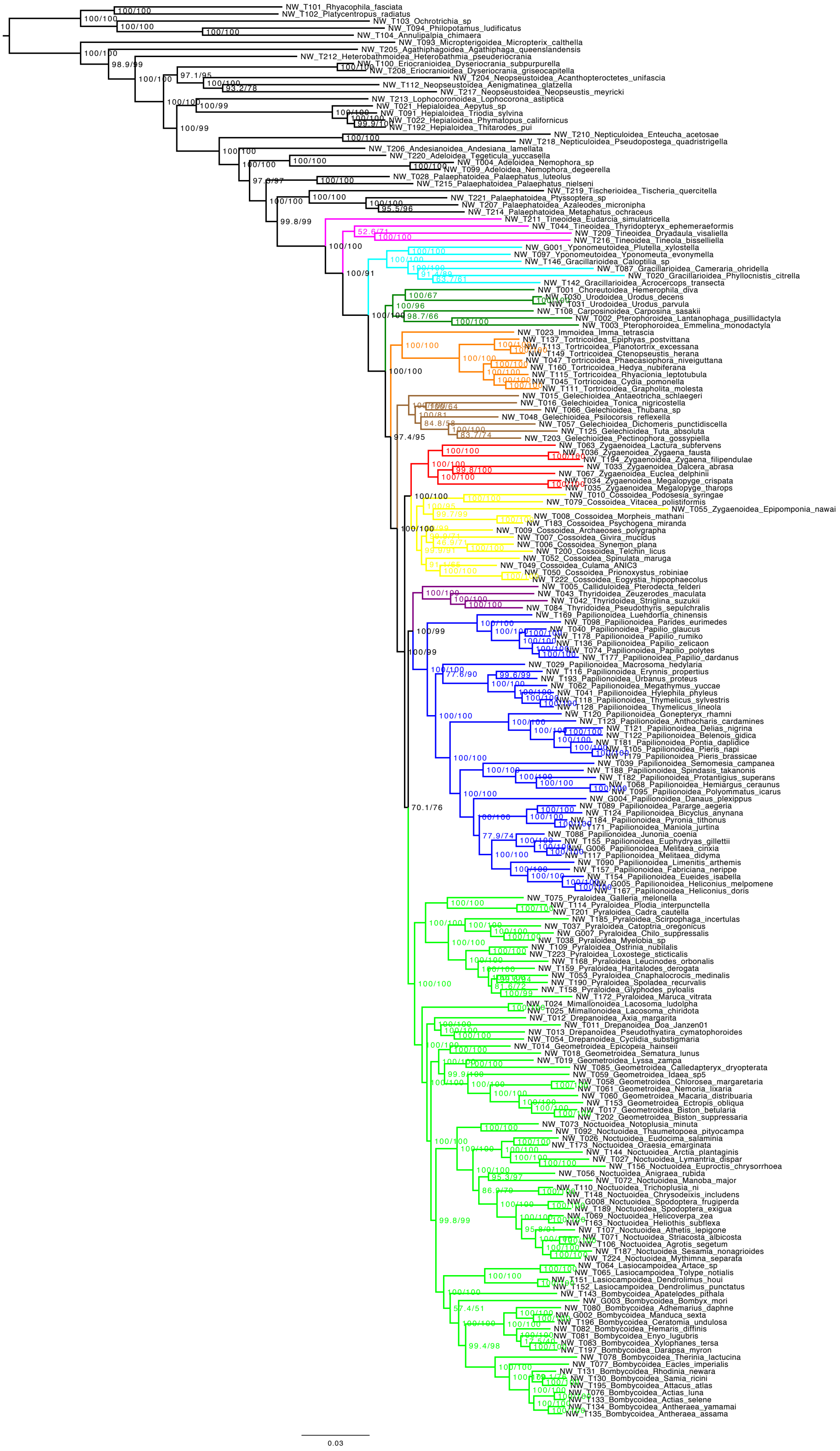

**Figure S1.** NT12_all.

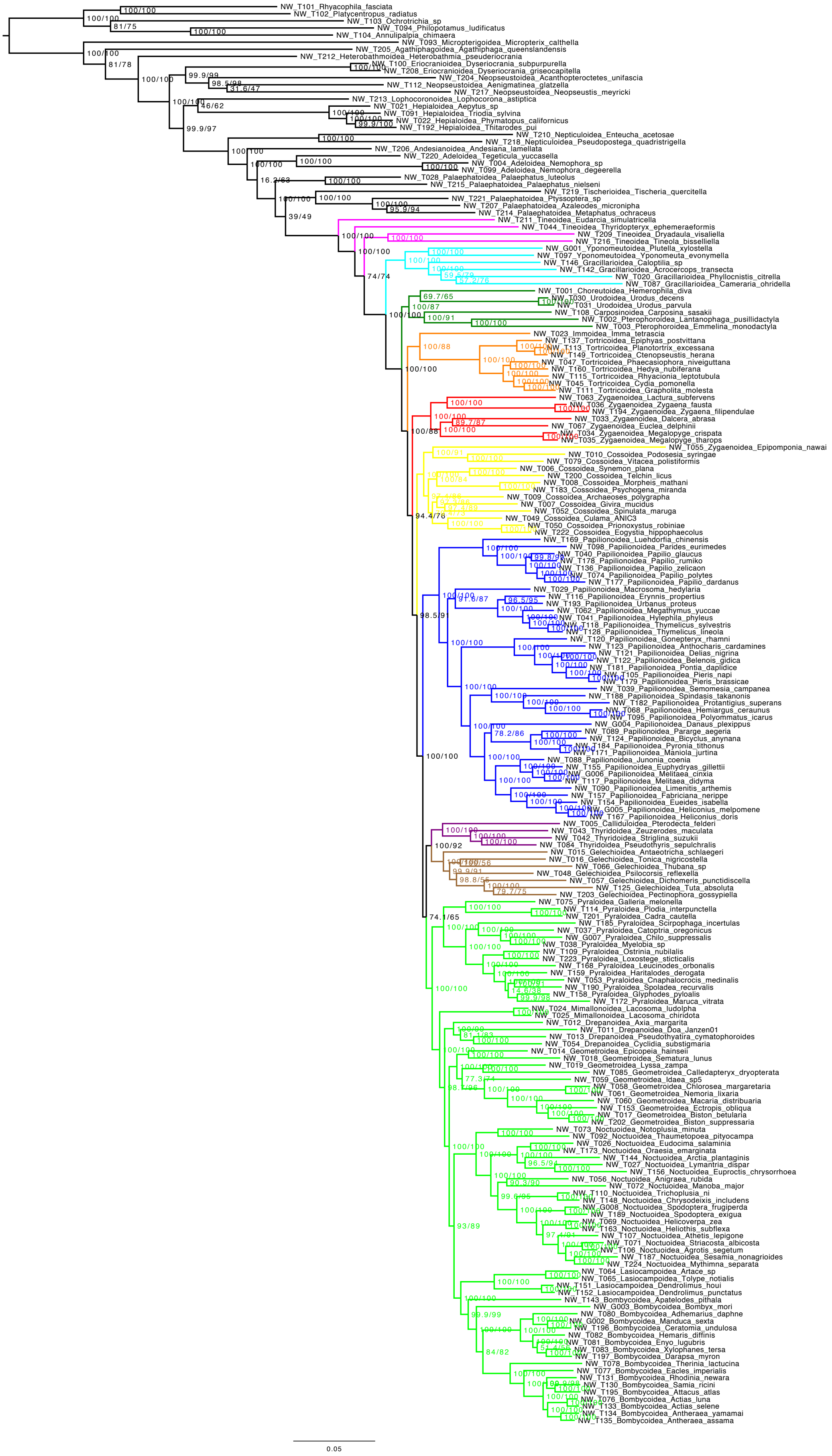

**Figure S2.** AA_all.

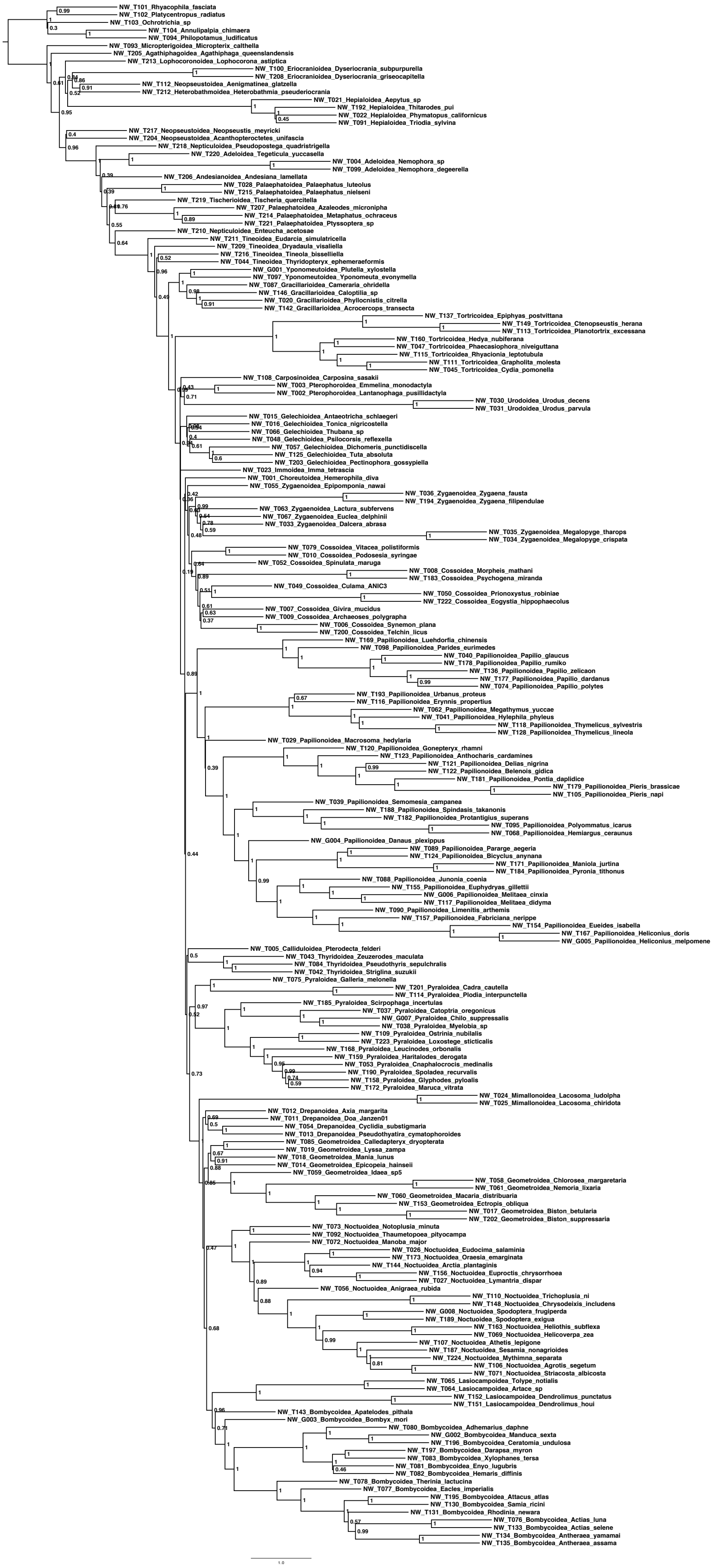

**Figure S3.** NT12 ASTRAL analysis. Numbers on branches represent local posterior probabilities as calculated in ASTRAL.

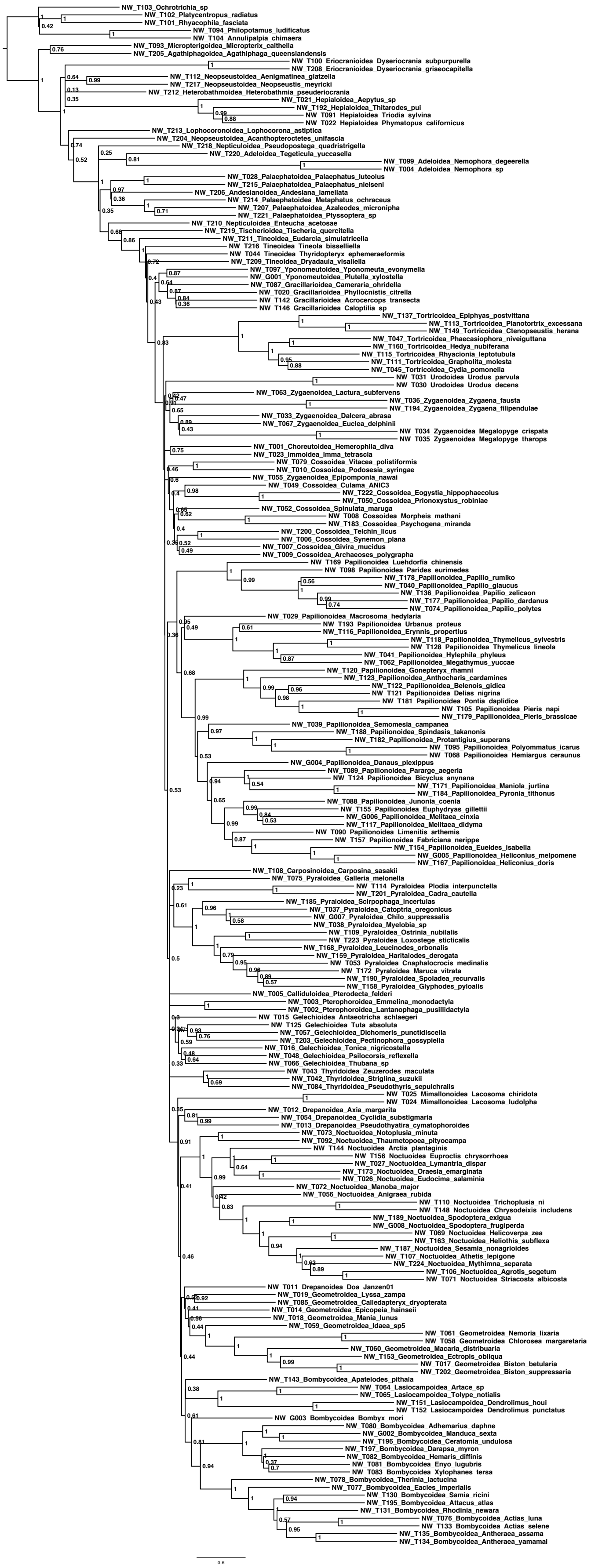

**Figure S4.** AA ASTRAL analysis. Numbers on branches represent local posterior probabilities as calculated in ASTRAL.

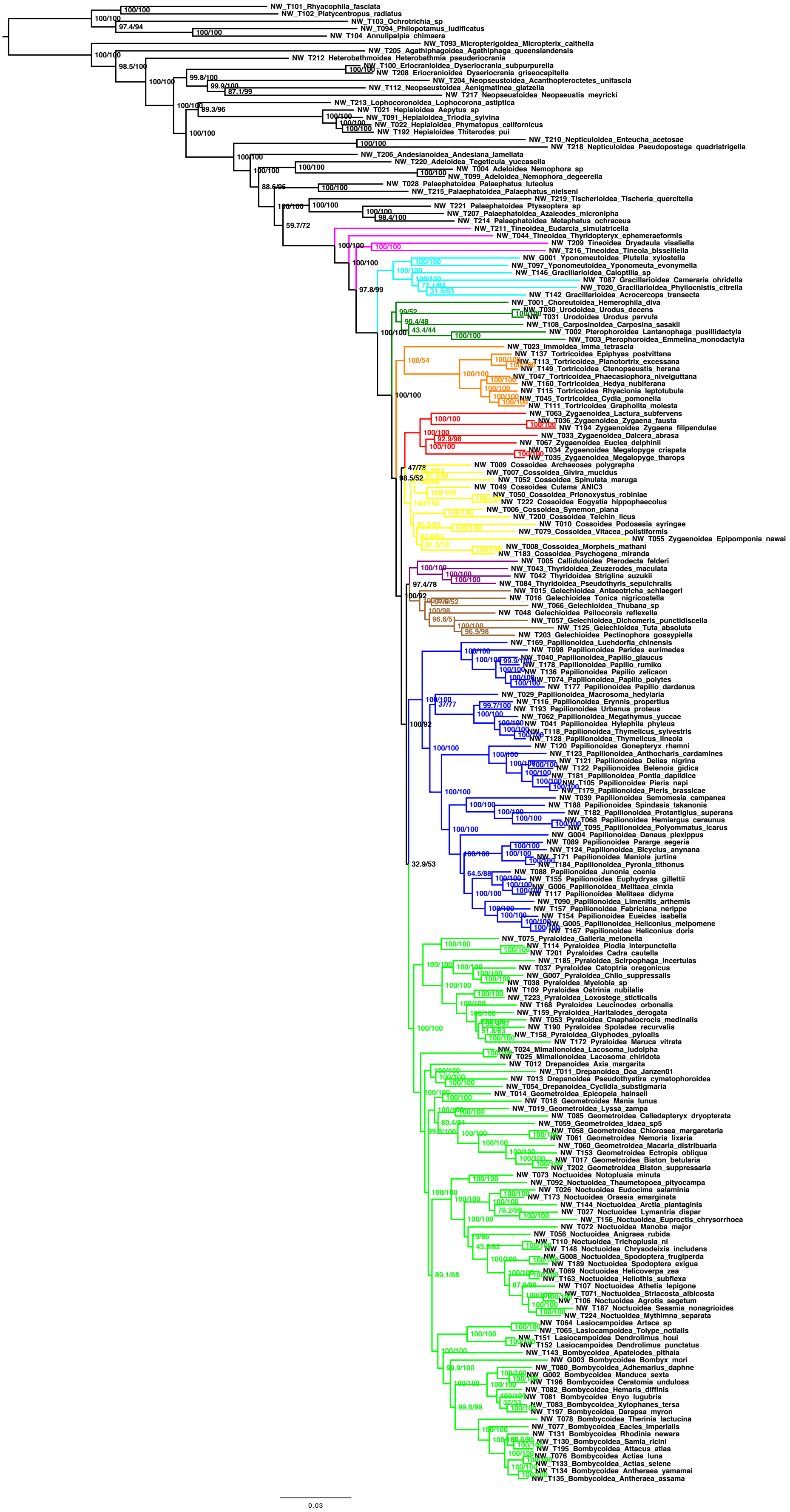

**Figure S5.** NT12_degen1.

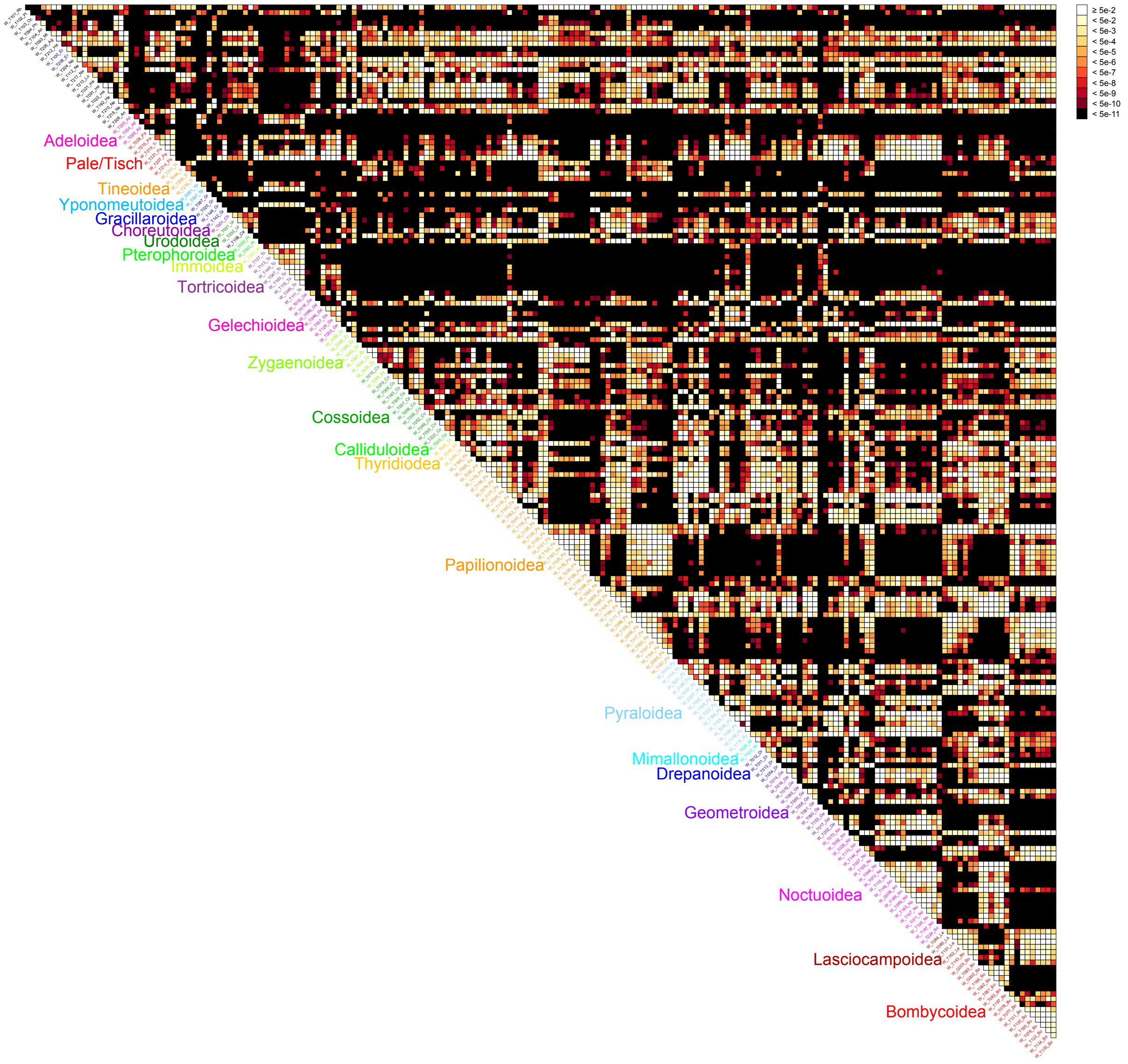


**Figure S6.** Bowker's test results for the NT12_all dataset.

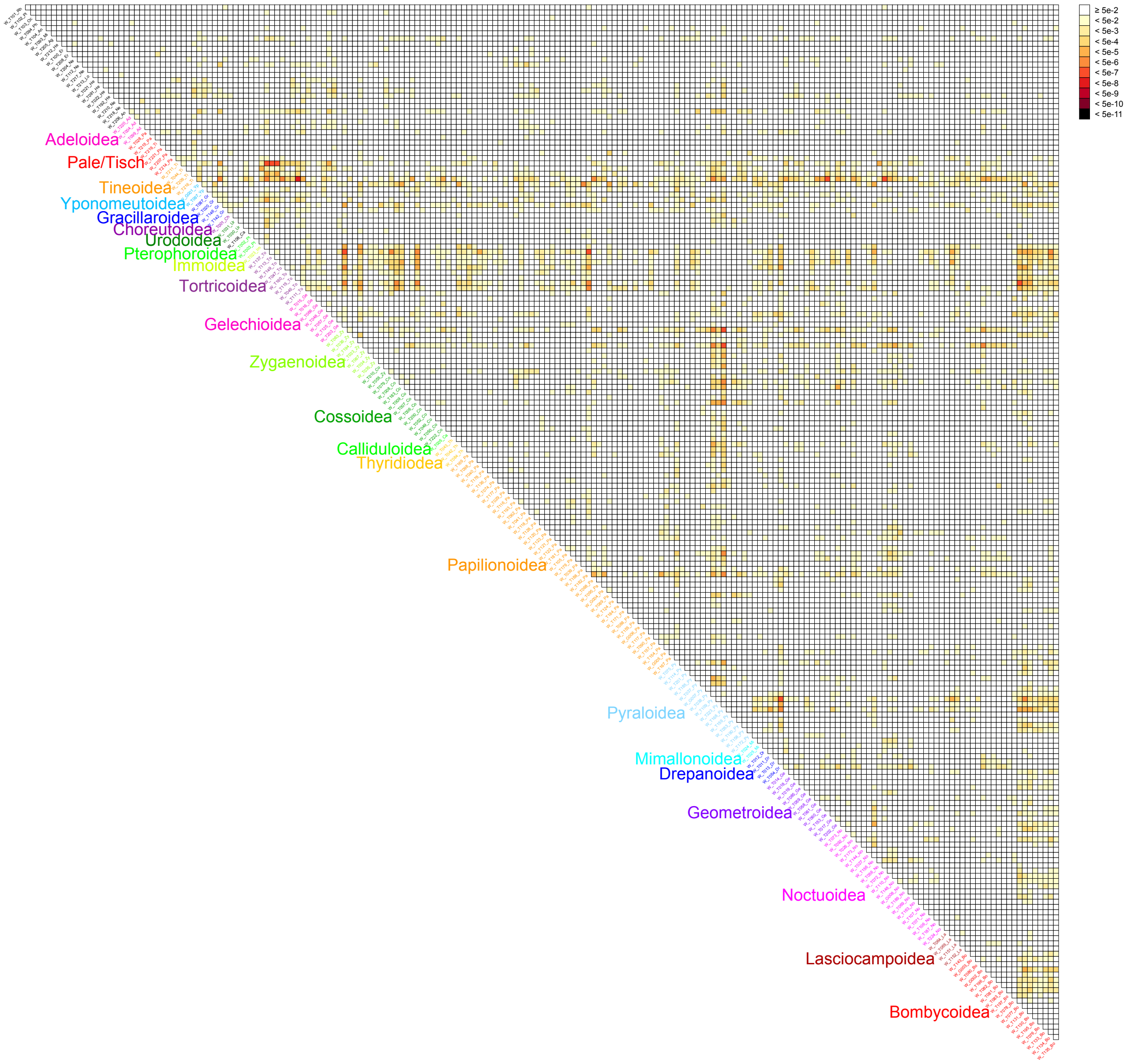


**Figure S7.** Bowker's test results for the AA_all dataset.

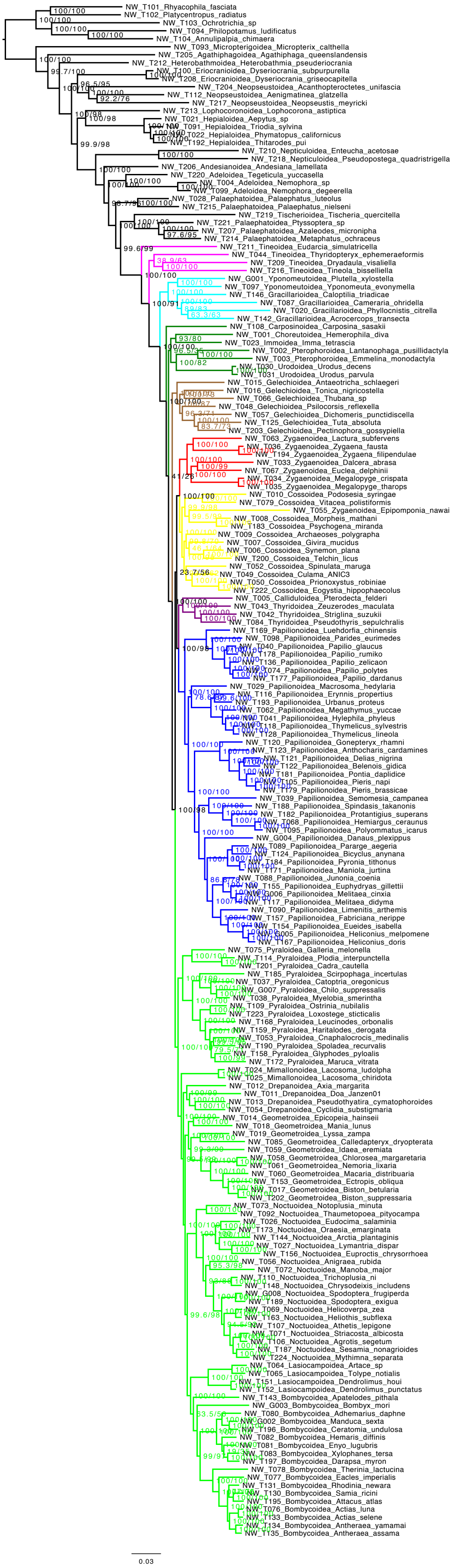

**Figure S8.** NT12_noTorts.

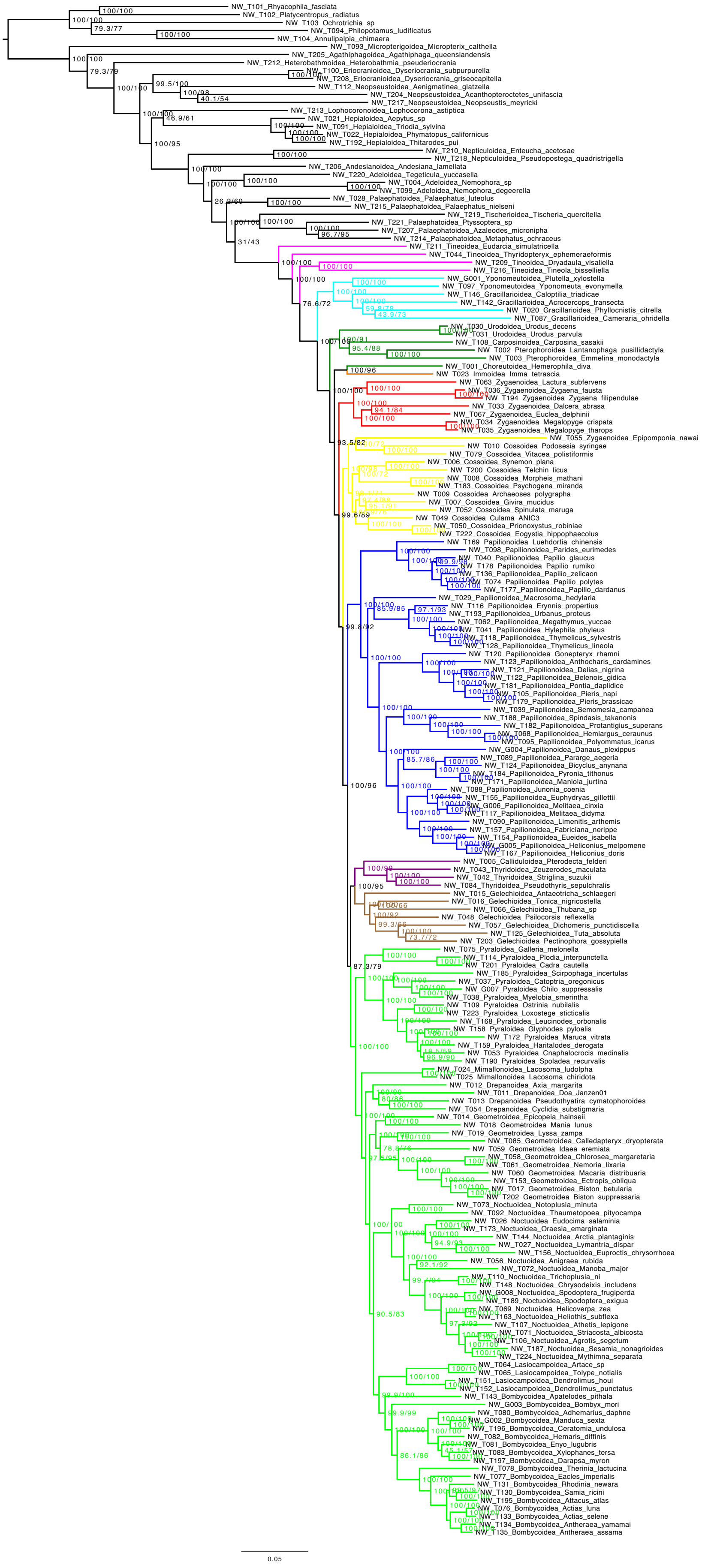


**Figure S9.** AA_noTorts.

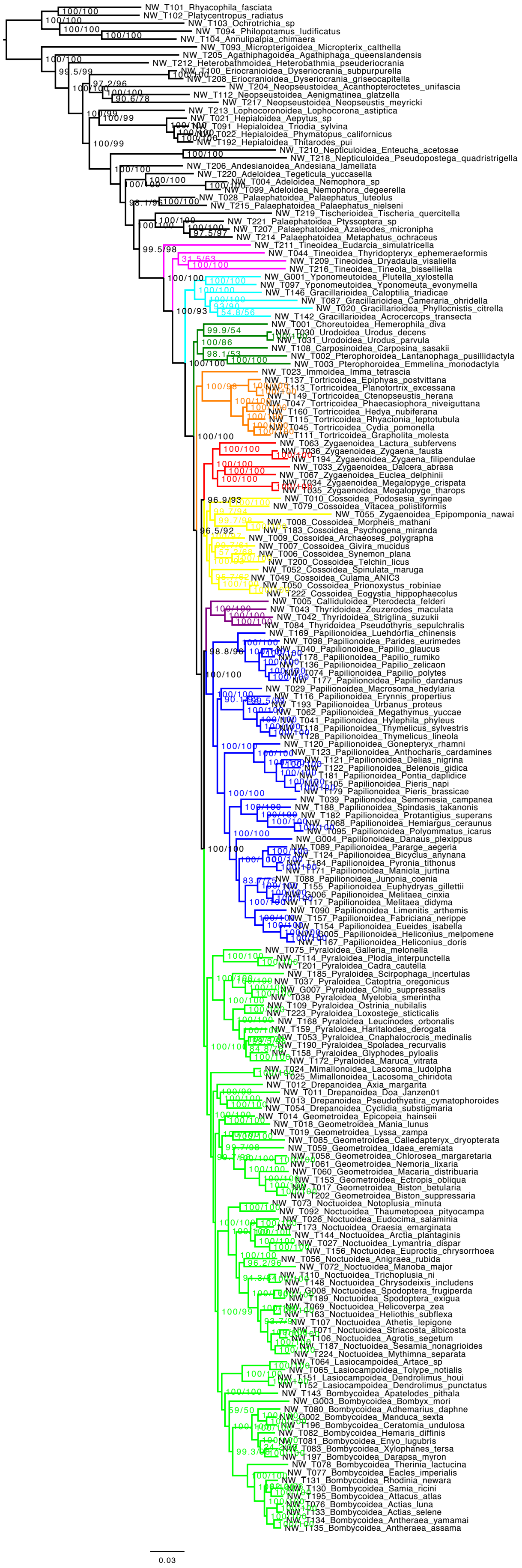

**Figure S10.** NT12_noGele.

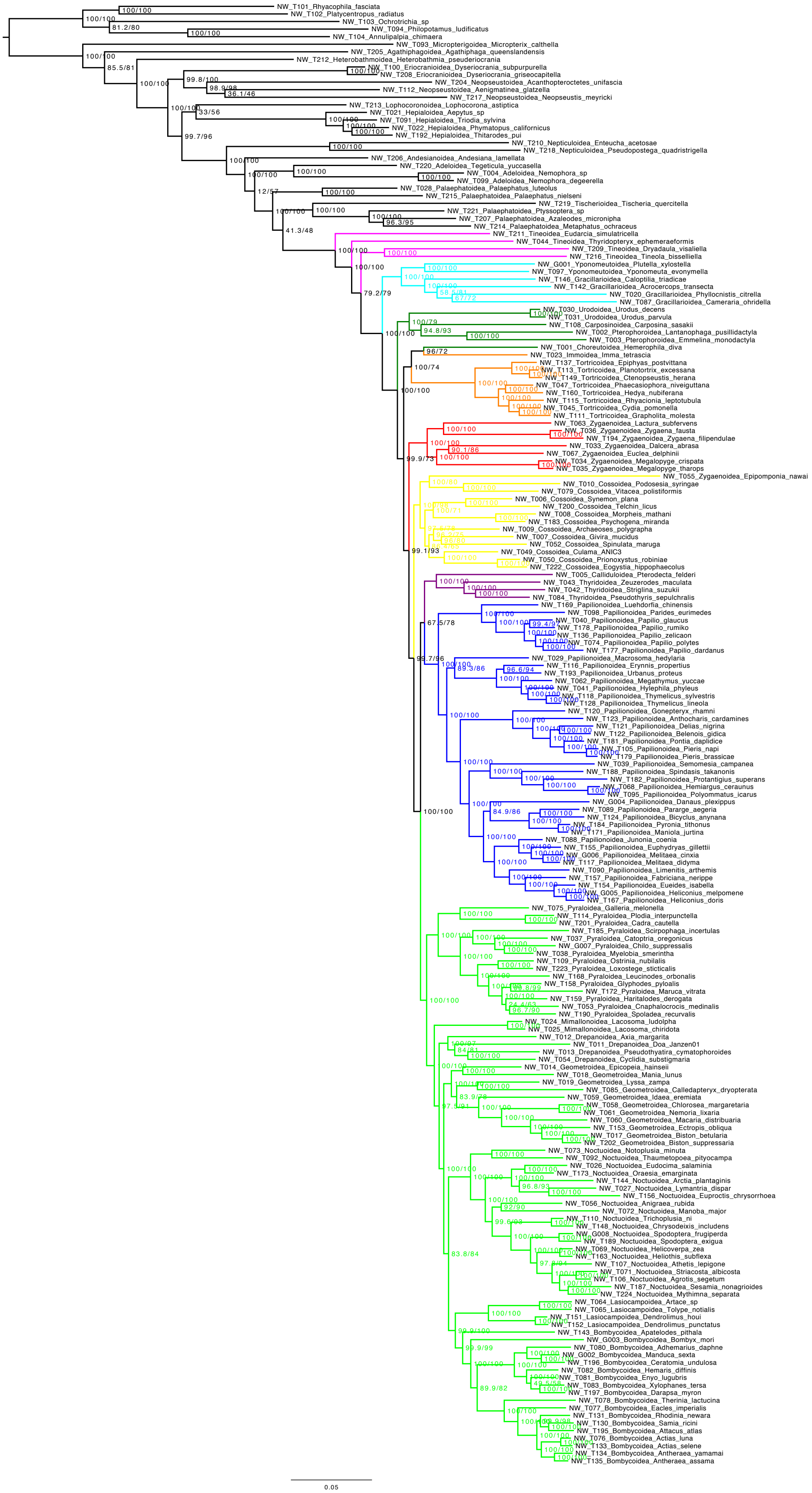

**Figure S11.** AA_noGele.

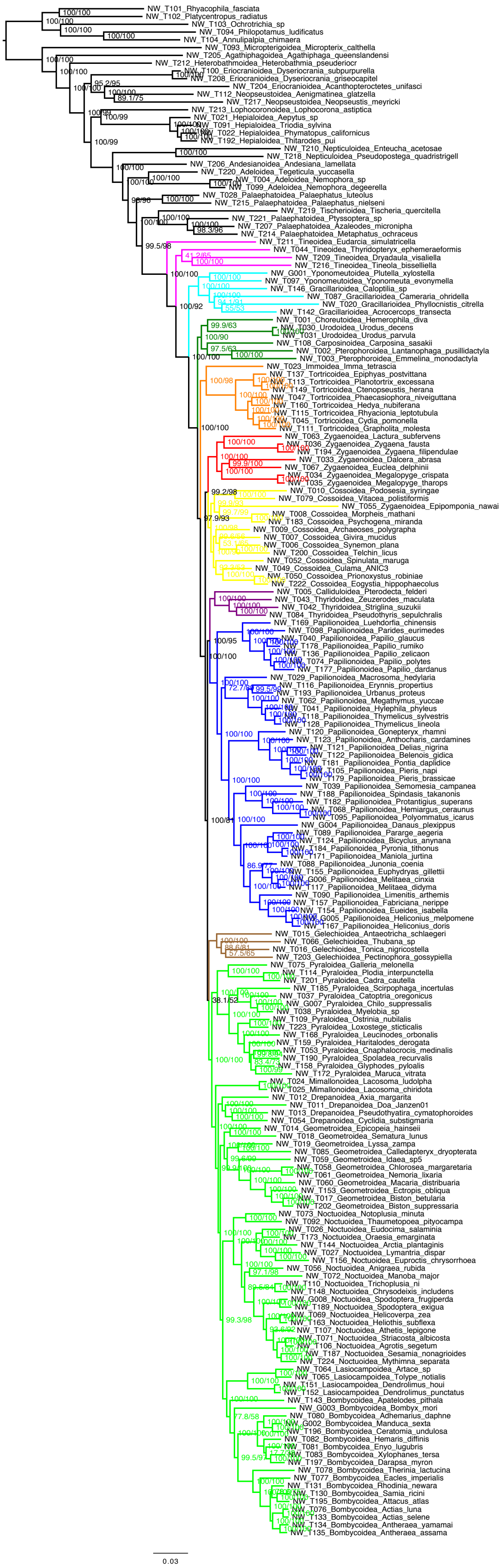

**Figure S12.** NT12_minus3Gele

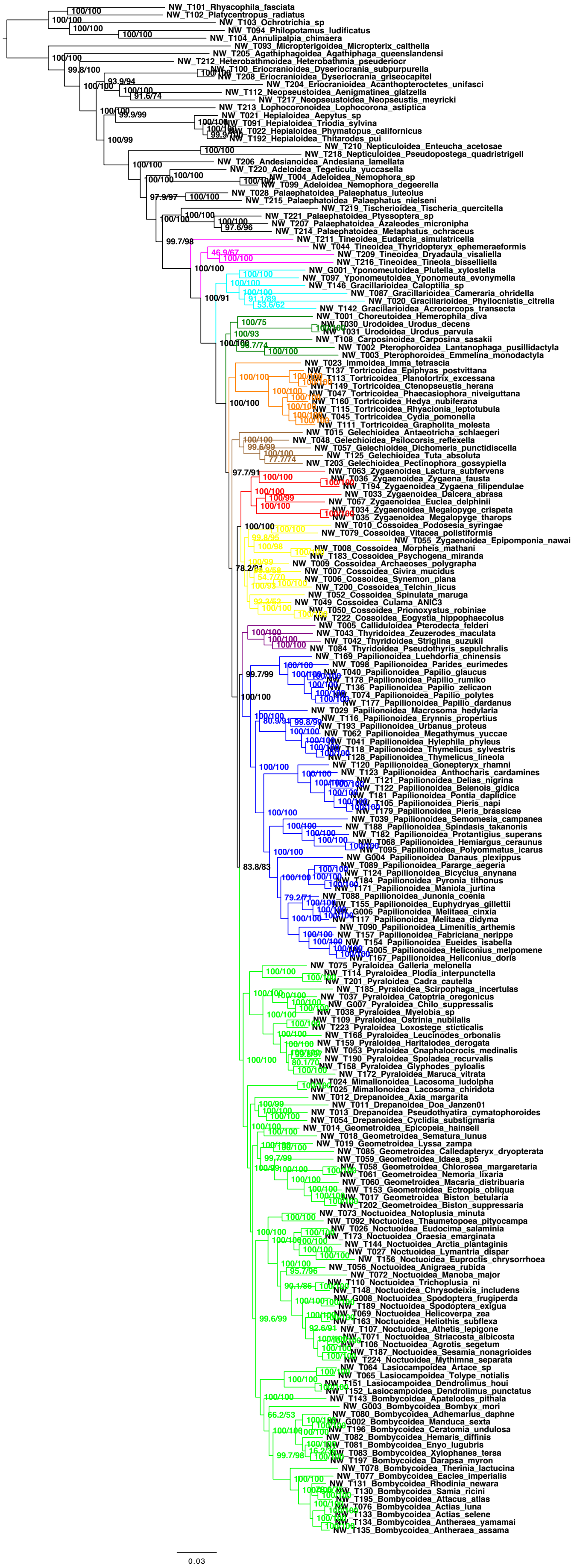


**Figure S13.** NT12_minus2Gele.

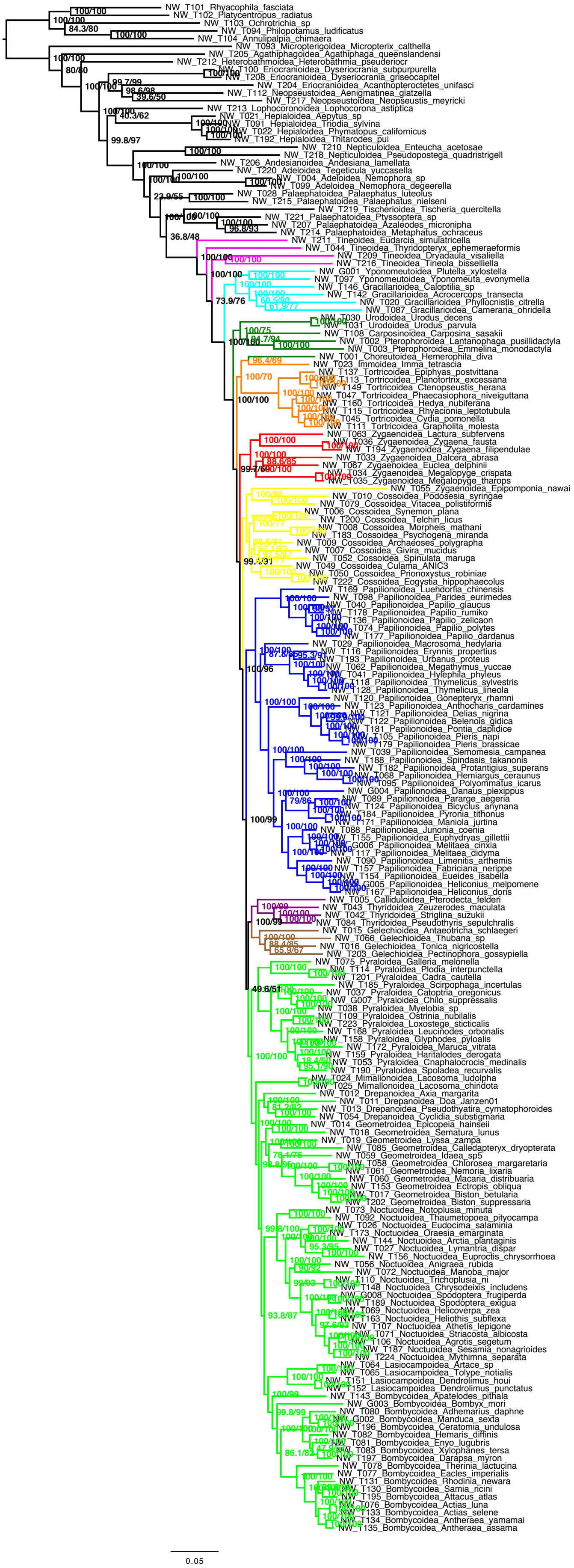

**Figure S14.** AA_minus3Gele.

**Rota et al.** The unresolved phylogenomic tree of butterflies and moths (Lepidoptera): assessing the potential causes and consequences

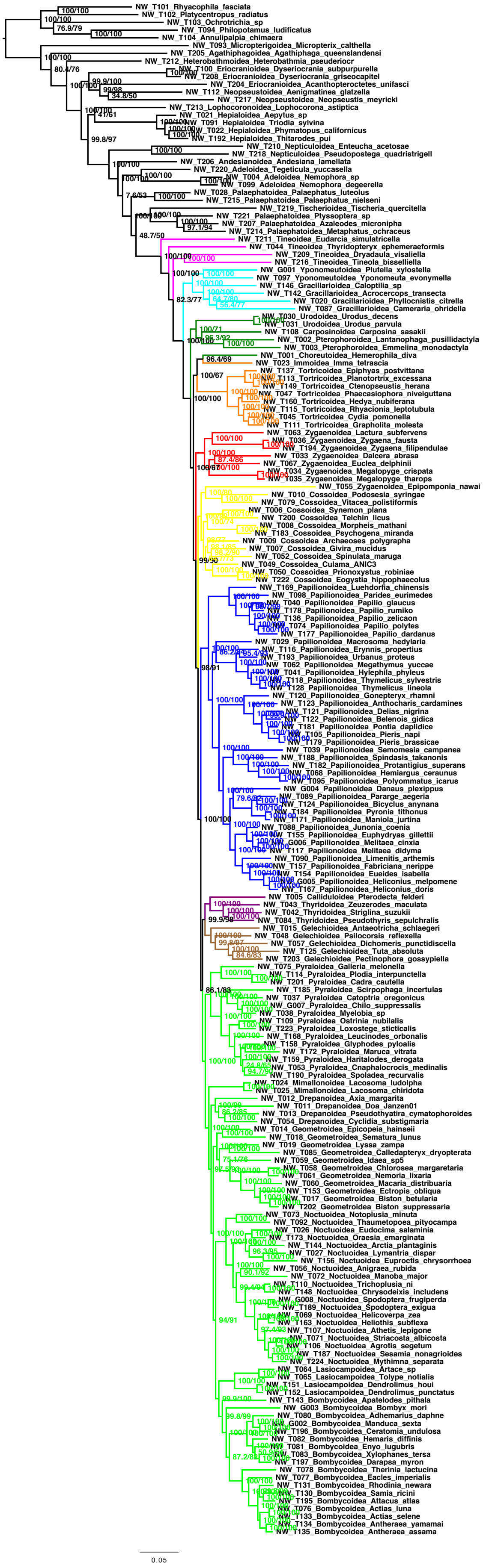

**Figure S15.** AA_minus2Gele.

**Rota et al.** The unresolved phylogenomic tree of butterflies and moths (Lepidoptera): assessing the potential causes and consequences

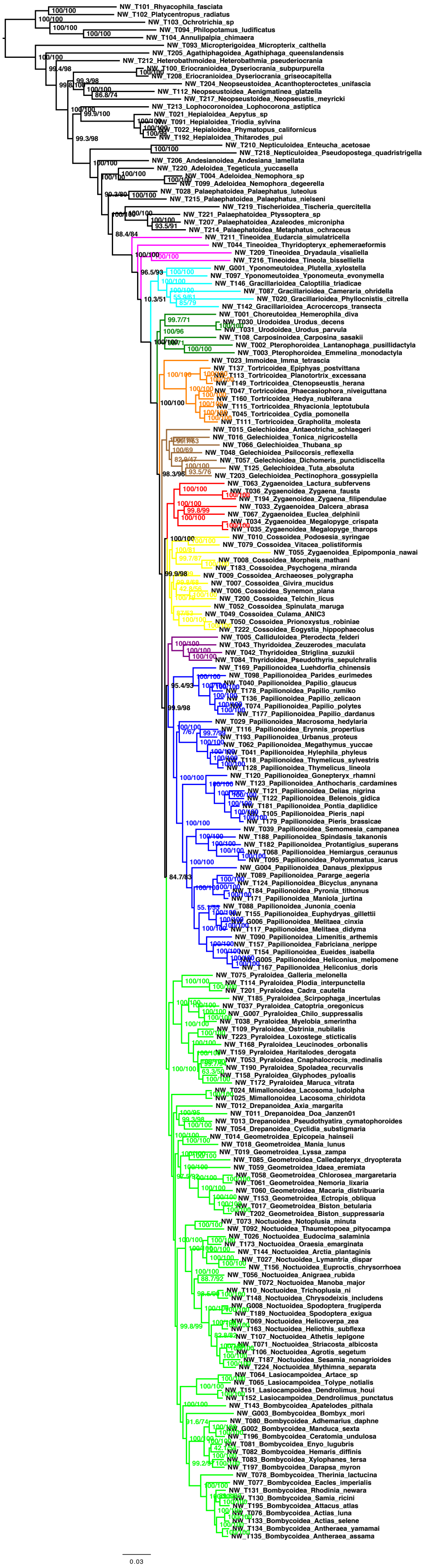

**Figure S16.** NT12_no_highGC.

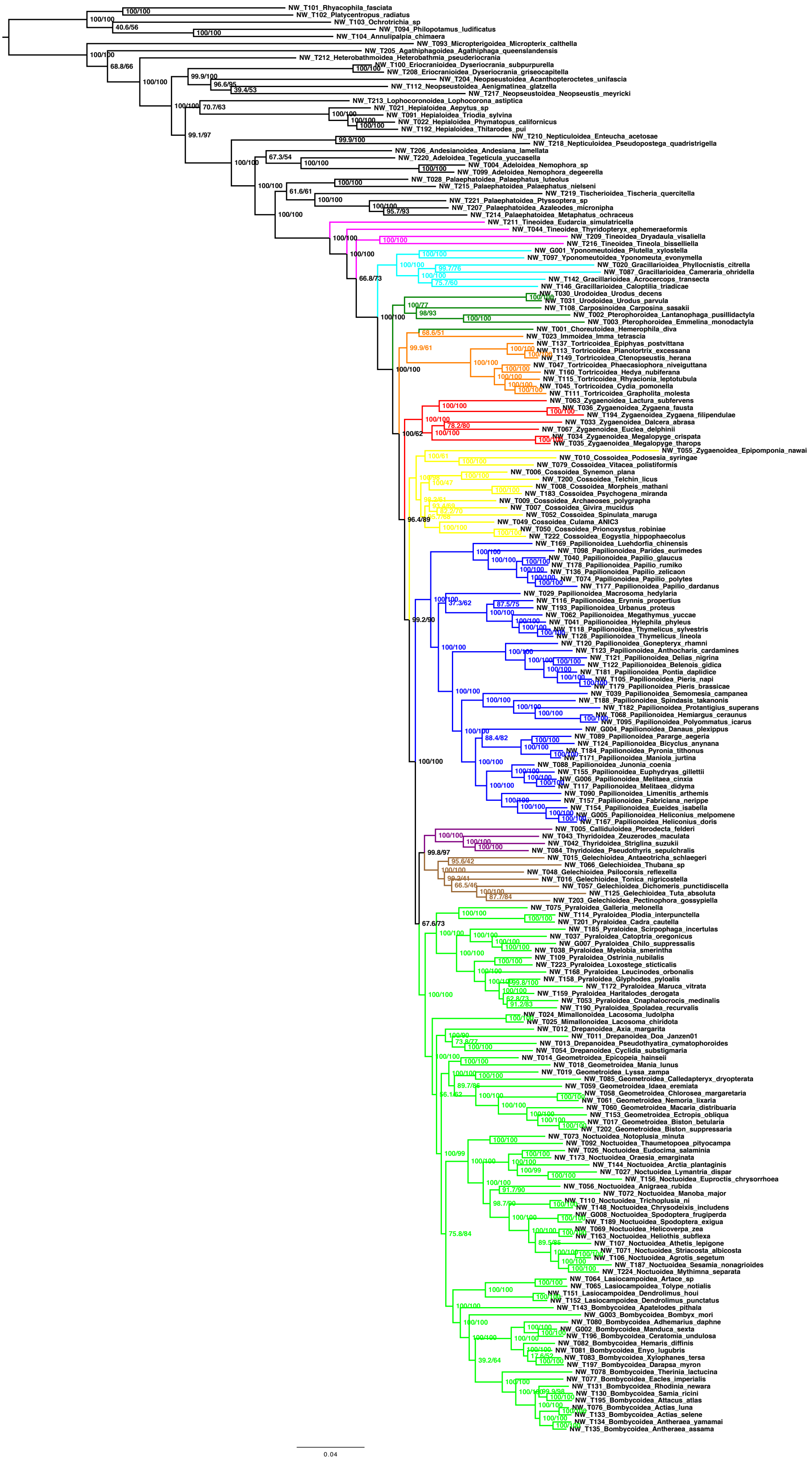

**Figure S17.** AA_no_highGC.

**Rota et al.** The unresolved phylogenomic tree of butterflies and moths (Lepidoptera): assessing the potential causes and consequences

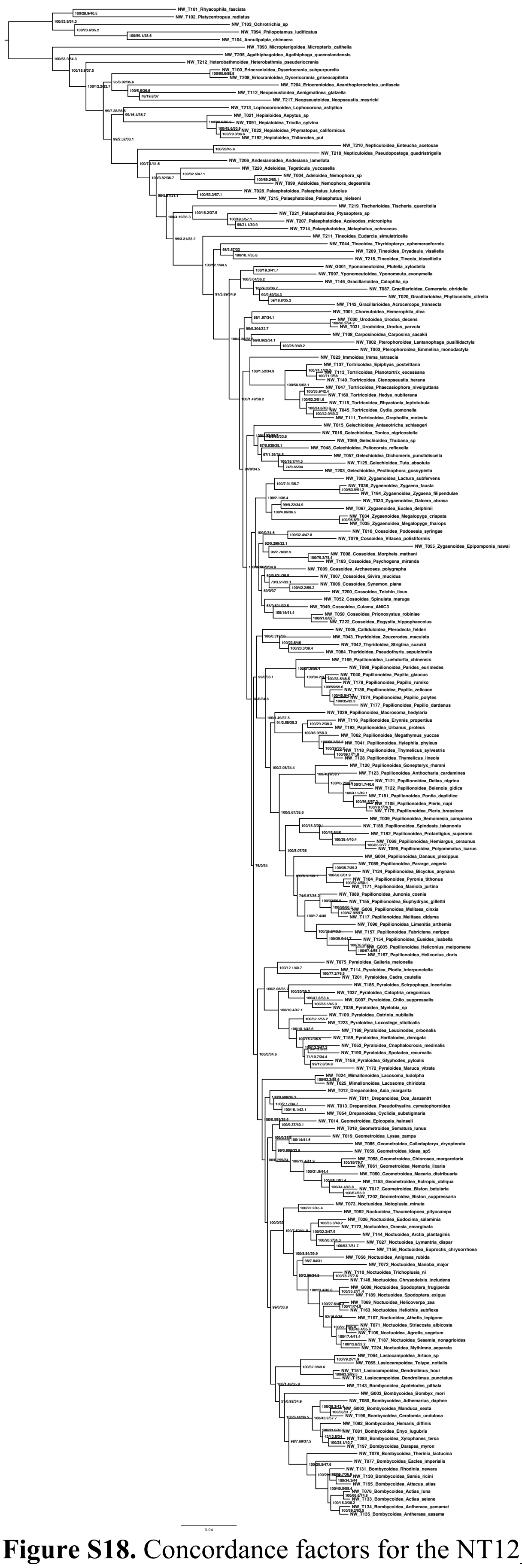

**Figure S18.** Concordance factors for the NT12_all topology. Numbers on branches are UFB/gCF/sCF.

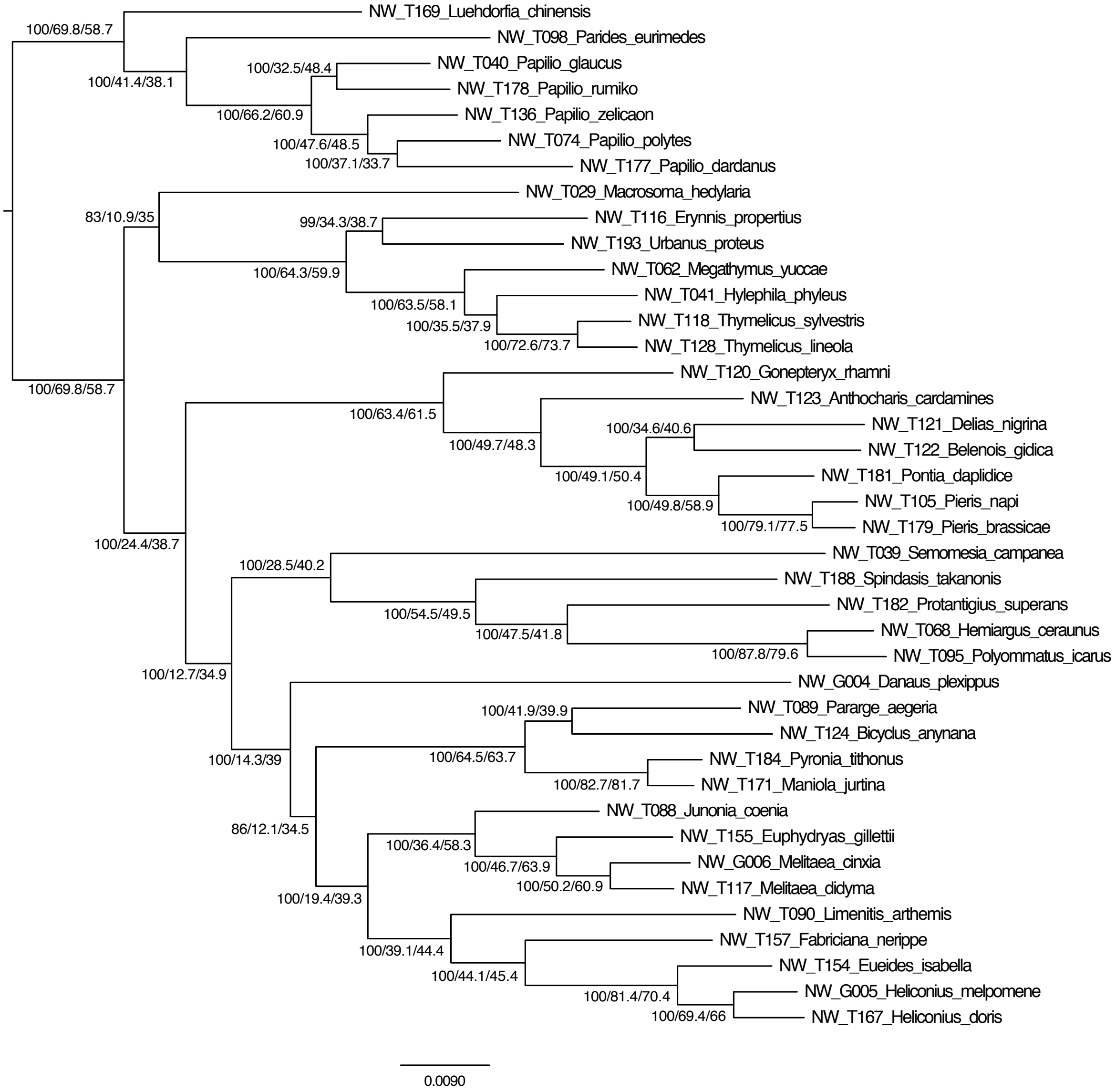


**Figure S19.** Concordance factors for the NT12 Papilionoidea topology. Numbers on branches are UFB/gCF/sCF.

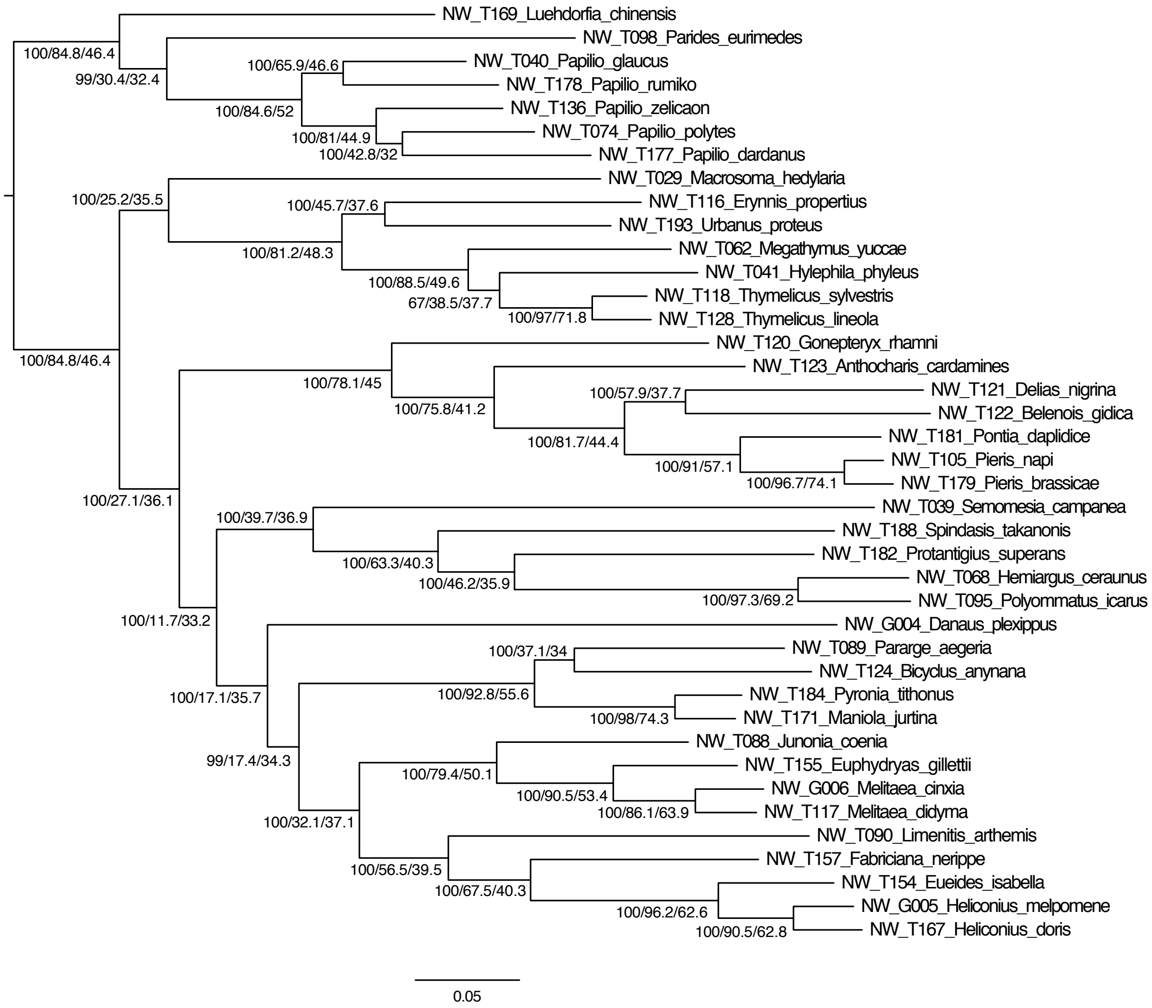


**Figure S20.** Concordance factors for the NT123 Papilionoidea topology. Numbers on branches are UFB/gCF/sCF.

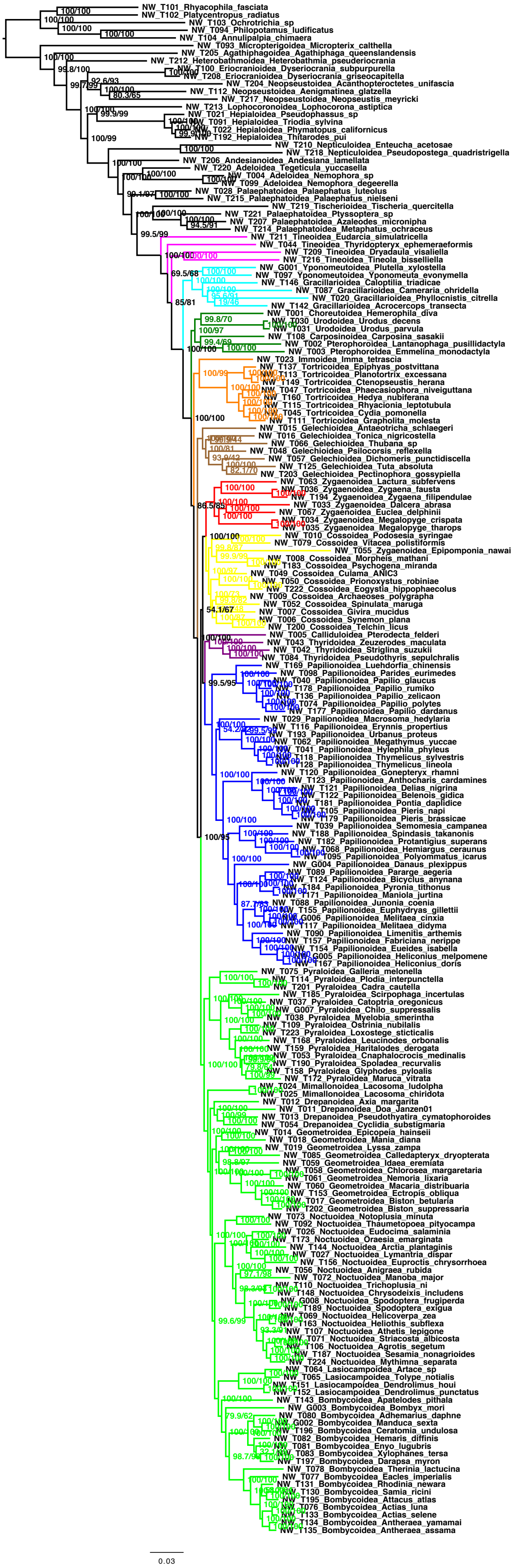


**Figure S21.** NT12_minus10.

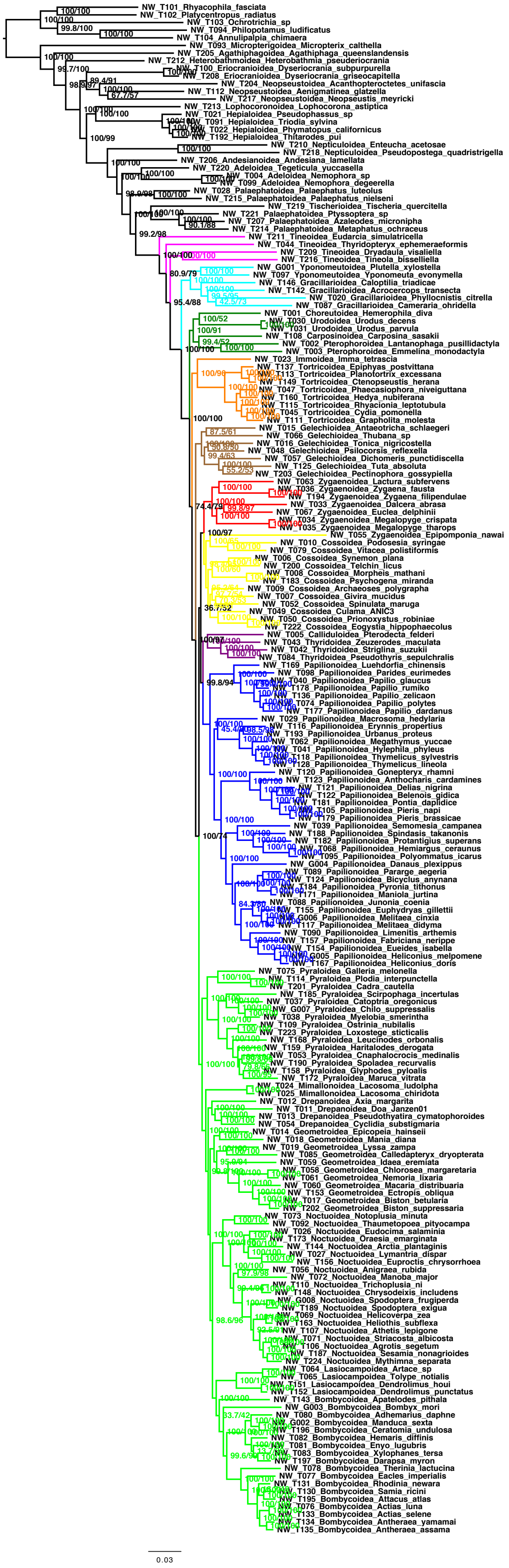

**Figure S22.** NT12_minus20.

**Rota et al.** The unresolved phylogenomic tree of butterflies and moths (Lepidoptera): assessing the potential causes and consequences

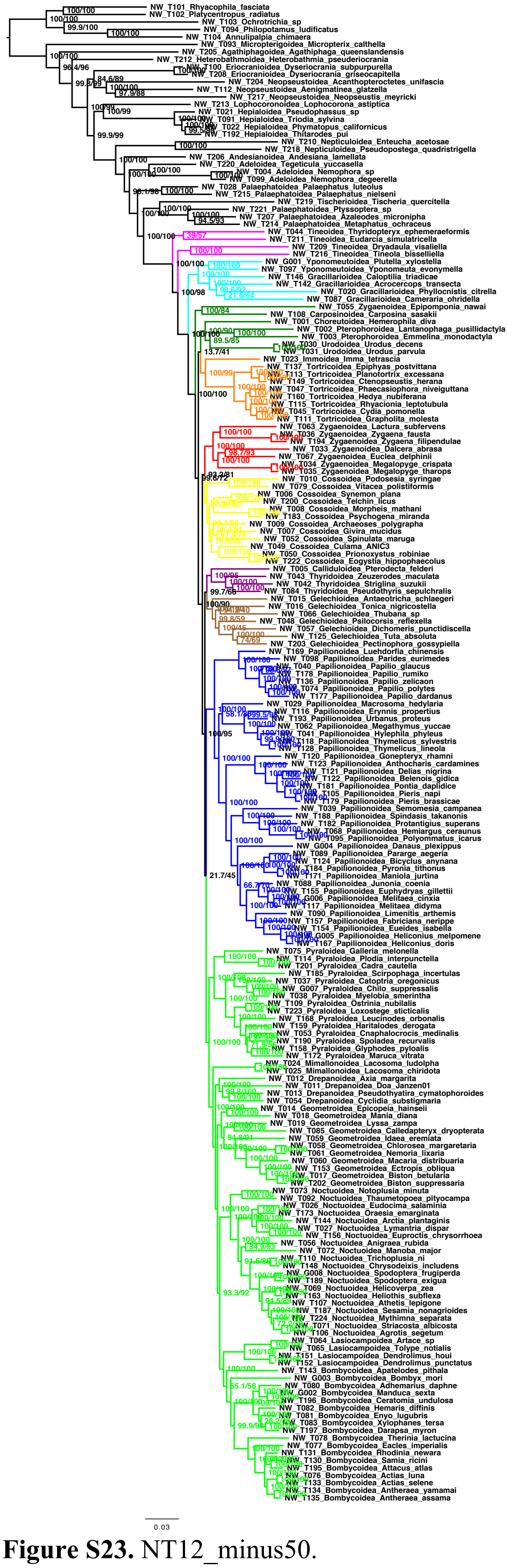

**Figure S23.** NT12_minus50.

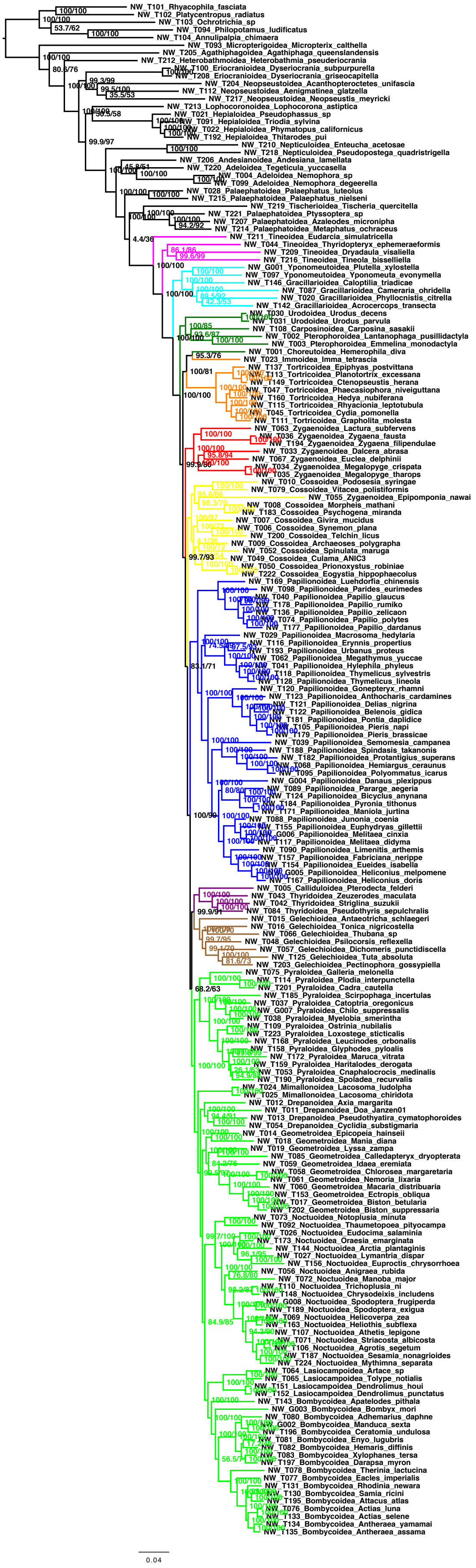

**Figure S24.** AA_minus10.

**Rota et al.** The unresolved phylogenomic tree of butterflies and moths (Lepidoptera): assessing the potential causes and consequences

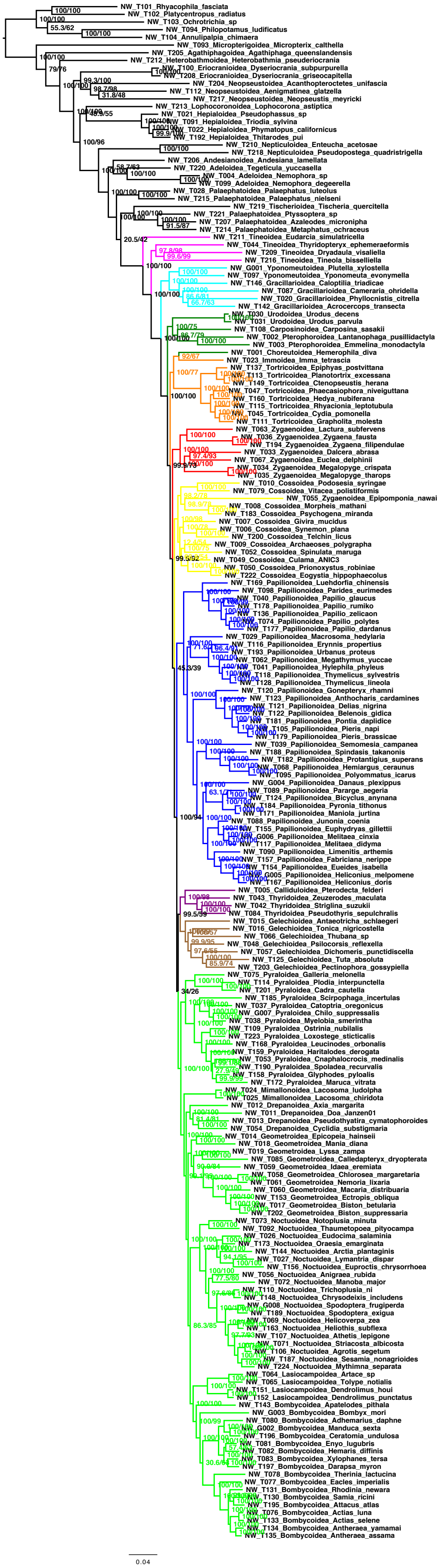

**Figure S25.** AA_minus20.

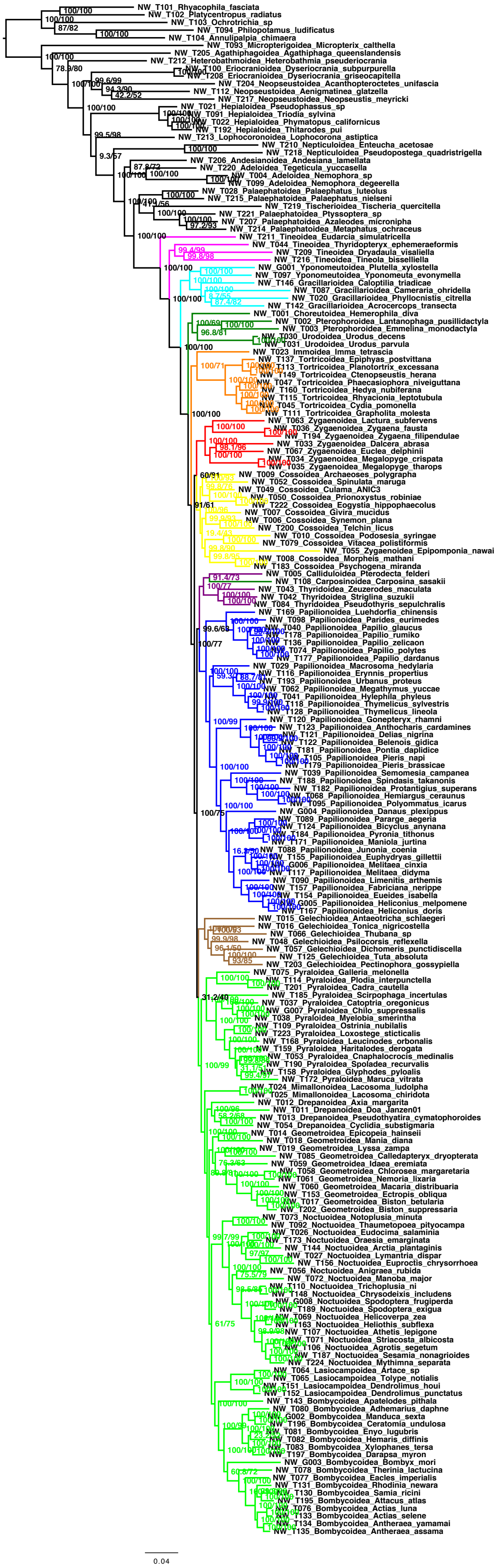

**Figure S26.** AA_minus50.
